# Autoencoder Model for Translating Omics Signatures

**DOI:** 10.1101/2023.06.08.544243

**Authors:** Nikolaos Meimetis, Krista M. Pullen, Daniel Y. Zhu, Avlant Nilsson, Trong Nghia Hoang, Sara Magliacane, Douglas A. Lauffenburger

## Abstract

The development of effective therapeutics and vaccines for human diseases requires a systematic understanding of human biology. While animal and in vitro culture models have successfully elucidated the molecular mechanisms of diseases in many studies, they yet fail to adequately recapitulate human biology as evidenced by the predominant likelihood of failure in clinical trials. To address this broadly important problem, we developed AutoTransOP, a neural network autoencoder framework to map omics profiles from designated species or cellular contexts into a global latent space, from which germane information can be mapped between different contexts. This approach performs as well or better than extant machine learning methods and can identify animal/culture-specific molecular features predictive of other contexts, without requiring homology matching. For an especially challenging test case, we successfully apply our framework to a set of inter-species vaccine serology studies, where no 1-1 mapping between human and non-human primate features exists.

## Introduction

Animal and cellular models are essential tools for studying the underlying biology of human diseases, but these insights are not always clinically translatable, resulting in the failure of numerous therapeutics in clinical trials^1,2^. A common approach is to choose orthologous biomolecules, including genes, proteins, and cellular pathways, to perform direct functional comparisons across species. However, functional divergence and the absence of orthologous biomarkers can hinder these direct comparisons between species ^3–5^. Furthermore, within the same species, the transcriptional response to chemical stimuli can be cell type-specific due to distinct genetic profiles, creating an additional barrier to understanding the mechanism of action of therapeutics^6–9^. Consequently, computational systems-based approaches are needed to gain a better understanding of the relationship between biological models and translate information gained from different model systems.

Advancements in sequencing technologies have enabled the generation of large-scale datasets from both animal and human species, facilitating more powerful analyses and comparisons of molecular features between different biological systems^2,3,10–13^. This has led to the development of numerous new statistical and machine learning models^3,13–17^ for identifying similarities between species and experimental models. Notably, most existing approaches focus on direct correlations between analogous biomarkers or processes across species despite known species and model system differences. In an attempt to address this challenge, Brubaker et al. proposed a technique called “Translatable Components Regression”^18^ (TransCompR), which maps human data into the principal component space of data from another species to identify translatable animal features that can predict human disease processes and phenotypes. Although this approach has been successfully applied to gain insights into some inflammatory pathologies^18,19^, it depends on homologs or comparable molecular features between species. While TransCompR is well suited to identify omics signatures in one species that is most germane for understanding phenotype characteristics in another, it is not centrally designed to integrate signatures across species. Moreover, this approach is by design only capable of deciphering linear relationships, thus potentially excluding non-linear biological relationships.

With the advent of deep learning, particularly autoencoders, there is great potential to develop approaches that can approximate the non-linear relationships underlying different biological systems and species. Autoencoders are artificial neural networks (ANNs) that can embed raw input data into a lower dimensional space from which the original data can be reconstructed ^20^. Autoencoders have been used in several tasks in biology including analyzing high dimensional data^21,22^, denoising single-cell RNA sequencing data^23–25^, deciphering the hierarchical structure of transcriptomic profiles^26,27^, and predicting gene expression caused by various stimuli^28–30^. One such model, DeepCellState^31^, focused on translating cellular states, can predict the transcriptional profile of a cell type after drug treatment based on the behavior of another cell type. However, similar to TransCompR, this approach depends on a 1-1 mapping of molecular features between cells to capture a global cell representation. Another recently proposed framework, is the compositional perturbation autoencoder (CPA)^32^. It can construct a basal latent space devoid of covariate and perturbation-specific signals, capturing only the basal cell state in single-cell RNA sequence data. CPA can be used to generate *in-silico* transcriptional profiles at the single-cell level for different perturbations, cells, and species, although it still requires mapping of orthologous genes. To overcome such limitations, an approach similar to those used in language translation autoencoder-based models, which create a global language representation^33,34^, may be useful and could aid biological inter-systems translation when 1-1 mappings between features do not exist.

In this study, we incorporate elements of the CPA approach with ideas from language translation models^33,34^ to develop an ANN framework hence referred to as AutoTransOP, Autoencoders for Translating Omics Profiles, which utilizes separate autoencoders for each biological system, enabling the mapping of samples into a global cross-model space, while providing feature importance estimates for various phenotype-prediction tasks. The basic model is trained to simultaneously minimize the reconstruction error of the input and the distance between samples coming from the same condition in the global latent space. Our framework is benchmarked, using the latest version of the L1000 dataset^12^, against the established approaches of TransCompR^18^, FIT^15^ and the ANN approach of DeepCellState^31^, which all require 1-1 feature mapping. We demonstrate that our approach outperforms FIT and DeepCellState, while there is no difference when comparing with TransCompR in cellular models. Additionally, we present several variations of the model and we illustrate the adaptability of our framework by applying it to data of varying omics type and sample size to answer different biological questions of interest. Furthermore, we demonstrate its biological interpretability, an aspect that deep learning models often struggle to attain, by using an integrated gradients approach ^35^. To analyze the performance of the model in inter-species translation we performed mouse^36^ to human^37^ translation of single-cell transcriptomics of lung fibrosis, as well as non-human primate^38^ to human translation^39^ of smaller-scale serology datasets to predict HIV vaccine efficacy in humans. The latter serves as a novel case study of cross-species translation where no 1-1 mapping between features exists. After building the model, we identified serological features in non-human primates that are predictive of protection against HIV in humans, without analogous features necessarily being present in human data. These findings demonstrate that features derived from this approach can be predictive of the phenotypic profile of another biological model without requiring them to be homologs, allowing us to maximize the amount of information we can capture from different model systems to advance our understanding of complex human disease biology.

## Results

### A flexible framework for omics translation

We developed a flexible artificial neural network framework (see methods) for omics translation across biological models. It consists of separate ANN encoders and decoders for each biological system, e.g. cell line or species, that share a global latent space (Figure 1a), eliminating the need for a 1-1 mapping between the features between systems. We implement two main variations of the global latent space intending to remove the system-specific effect of perturbations. The first variation, which is also the main model variation of this framework (model variation 1), consists of a single global latent space that is created by maximizing the similarity of embeddings derived from the same condition/perturbation in a different species or cell line. The second variation (model variation 2) is based on the recently published compositional perturbation autoencoder (CPA)^32^, where there are two separate latent spaces: 1) a global/basal latent space and 2) a composed latent space. The global latent space expands on the first variation with an additional discriminator that attempts to remove the cell-line or species effect by penalizing models where the classifier can detect from which encoder the latent representation originates^32^. In the composed latent space, a cell/species classifier is simultaneously trained to ensure there is a cell/species effect, which is either added through a trainable covariate vector^32^ or added through two intermediate ANNs, allowing for non-linearity. We utilize integrated gradients^35^ to estimate feature importance for various predictive tasks. Lastly, we also introduce a variation (model variation 3), with one single global latent space, where a classifier is simultaneously trained on the global latent space (see methods). This is a contradictory learning task where the framework attempts to simultaneously remove the cell line or species effect globally but also hides cell or species information in a few of the latent variables.

**Figure 1:**
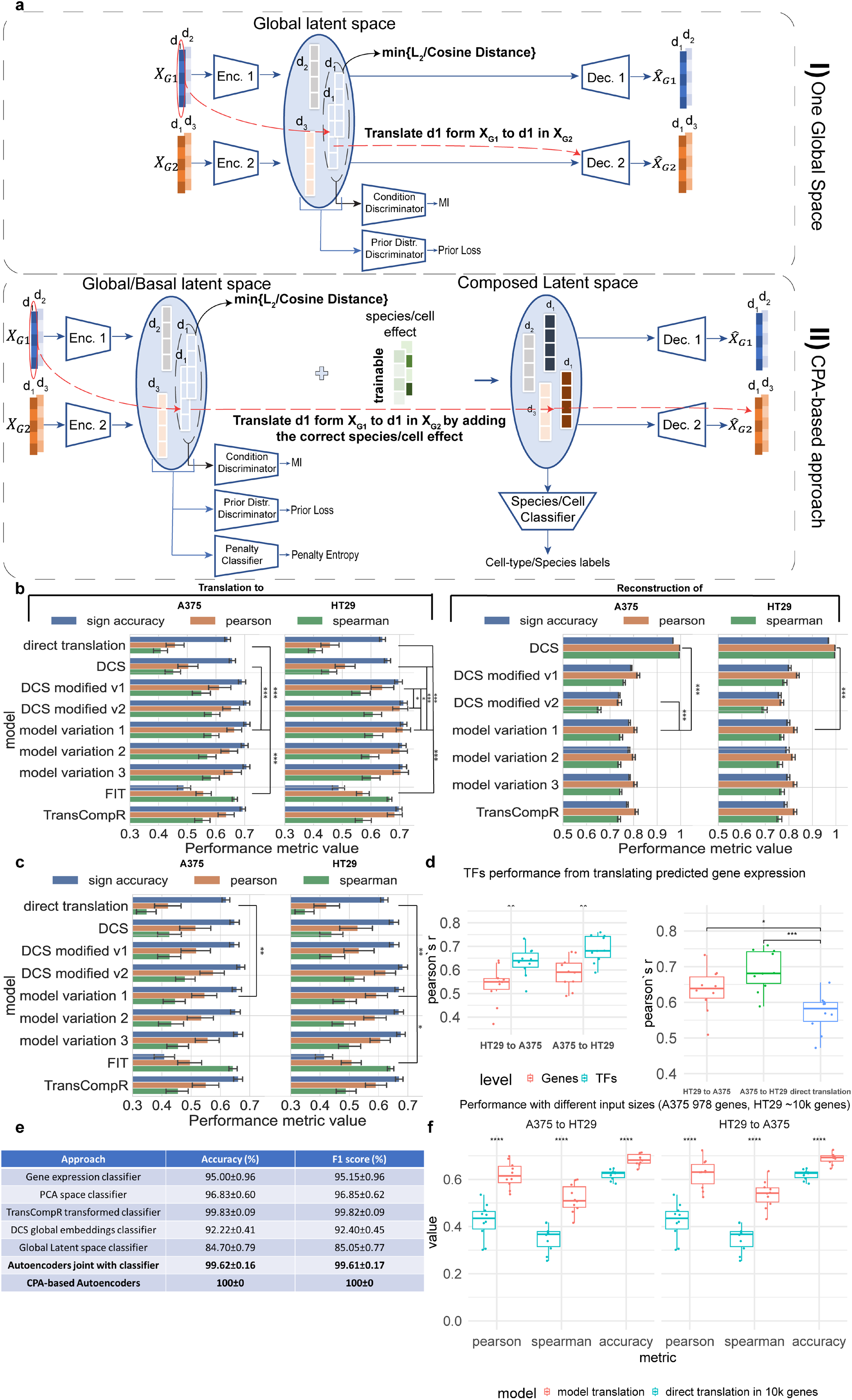
Model architecture and basic performance metrics. **a)** Framework architecture main variations: I) **model variation 1:** One global space is constructed by mapping omic profiles in a space where the distance between embeddings coming from the same perturbation is minimized. II) **model variation 2:** CPA-based architecture where the latent space is separated into two, one global devoid of species/cell effect and a composed latent space. **b)** Model performance in reconstructing and translating gene expression profiles between the two cell lines with the most common perturbations in the L1000 dataset, A375 and HT29, by using only the 978 measured landmark genes. The **model variation 3** is the one with a classifier simultaneously trained in one global latent space. For DCS modified v1-v2 see Supplementary Methods 2.2-2.3. It is worth noting that DCS modified v2 has a distance term and a direct translation term in its training loss. **c)** Model performance in reconstructing and translating gene expression profiles between A375 and HT29 by using all 10,086 genes that are either measured or belong to those that are well-inferred computationally. **d)** Performance in inferring transcription factors activity by using the translated/predicted gene expression. **e)** Performance in correctly classifying cell lines in different cases. **f)** Performance by using different inputs in the L1000.

### Benchmarking reconstruction and translation of gene expression profiles between two cell-lines

First, we compared our ANN framework with state-of-the-art techniques in the context of translating homologous genes between in-vitro models within the same species. We use the L1000 transcriptomics dataset^12^ to benchmark different approaches to translate the effects of perturbations between different human cell lines. The two main variations of our approach, as well as the variation where a classifier is simultaneously trained on the global latent space, are compared with three previously published approaches (DeepCellState^31^, FIT^15^, TransCompR^18^). As a baseline, the models are also compared to “direct translation”, i.e. directly using the gene expression profile in one cell line as a prediction for the effect in another cell line. We evaluate the models both on the task of translating the gene expression profile between cell lines, as well as the task of accurately reconstructing the gene expression for the same cell line. We evaluate them using several different metrics: i) Pearson’s correlation between predicted gene expression and actual gene expression, ii) the per sample Spearman’s rank correlation, and iii) the accuracy in correctly predicting the sign of drug-induced gene expression.

When utilizing the 978 landmark genes measured in the L1000, all of our framework’s variations provide a statistically significant increase in performance compared to the direct translation across all metrics (Figure 1b, Supplementary Tables 1-4 with p-values). When translating from the HT29 cell line to A375, our main model variation outperformed FIT^15^ and the basic DeepCellState^31^ (DCS) methods. When translating in the reverse direction, from A375 to HT29, our framework also outperforms the different modifications of DCS (Figure 1b). It can be noted that the 2^nd^ modification of DCS that enforces similarity in the latent space like our model, also outperforms the basic DCS, which may support the importance of enforcing similarity in the global latent space via some distance metric. For reconstruction of the input within a single cell line, the basic DCS approach outperforms the other approaches, at the expense of its translation performance. On this metric, our approach performs well and comparably with the other methods (Figure 1b). The alternative variations of our framework also perform comparably well.

When using the L1000 dataset with computationally imputed expression of 10,086 genes, the performance of all approaches drops, though still better than the baseline. There is generally no statistically significant difference between our approach and the other state-of-the-art approaches (Figure 1c). Interestingly, our approach performs better than direct translation also in the case of using different genes as input for each cell line, e.g. using only the 978 landmark genes for the A375 cell line and all the 10,086 genes for HT29 (Figure 1f). The performance is comparable to that using the same genes for both cell lines, indicating the potential to later extend the method in cases where no 1-1 mapping exists.

### Performance in using predicted gene expression to infer transcription factor activity

While the performance was worse in predicting the full set of 10,086 imputed genes, we reasoned that these imputed transcriptomic profiles may still be useful as input into different aggregation methods, e.g. to infer the activity of transcription factors (TFs). When we inferred transcription factor activity (see methods), model performance increased relative to using all 10,086 genes and was comparable to that in the case of the landmark genes (Figure 1b and 1d). Our model was not as successful at predicting gene set enrichment (Supplementary Method 1, Supplementary Figure 3). Autoencoders have been previously shown to be capable of capturing regulatory relationships between genes^26,31^ but, to our knowledge, not gene set enrichment, which might explain why we observed increased performance only when inferring TF activity.

### Creating cell-line-specific regions in the latent space enables robust cell classification

It is important to evaluate whether the cell line or species effect is successfully added to the composed latent space and whether the framework can retrieve it. To establish the ability of the model to capture cell-line-specific information we evaluated the performance in classifying the cell line when using all 10,086 genes (Figure 1e) and the landmark genes (Supplementary Table 11) of the L1000 dataset. The performance of ANN classifiers trained directly on the L1000 gene expression data serves as the baseline. Classifiers built with pre-trained embeddings, from DCS or our framework with one global latent space, are expected to have lower performance than the baseline as these approaches generate embeddings aiming to filter the cell-line effect as much as possible. Our framework seems to be better at “forgetting” the cell line of origin in the global space than DCS, thus generating more global embeddings (Figure 1e). Interestingly, when simultaneously training a classifier in the global latent space we can outperform the baseline while the cell-line effect is still partially filtered in the higher dimensions (Supplementary Figure 4). The CPA-based model in the composed latent space classifies cell lines with 100% accuracy, even though the similarity of input gene expression data between training and validation sets, as well as the latent space embedding similarity, is generally low (Supplementary Figure The CPA-based framework can create very well-separated cell-line-specific regions (Supplementary Figure 6) in the composed latent space, indicating the framework’s ability to shape the latent spaces with robust cell-line-specific regions and explaining the observed accuracy.

### Analysis of the framework’s dependence on different aspects of the data

We further investigated how the performance of the framework was influenced by different factors, focusing on the CPA-like approach. The framework has similar behavior and performance across cell-line pairs (Figure 2a). For all cell lines, ∼600 total training samples are sufficient to train a high-performance model. Some cell-line pairs perform slightly worse, as the original correlation between the same perturbations in the cell-line pair correlates with the model’s performance (Figure 2b). Interestingly, the amount of paired conditions, meaning similar perturbations across biological systems, required to successfully facilitate translation can be as low as ∼10-15% of the samples being paired (Figure 2c). Finally, it seems the model is not affected by a moderate imbalance in the number of conditions coming from each cell line (Figure 2d). Similar trends are observed when using 10,086 genes (Supplementary Figure 7).

**Figure 2:**
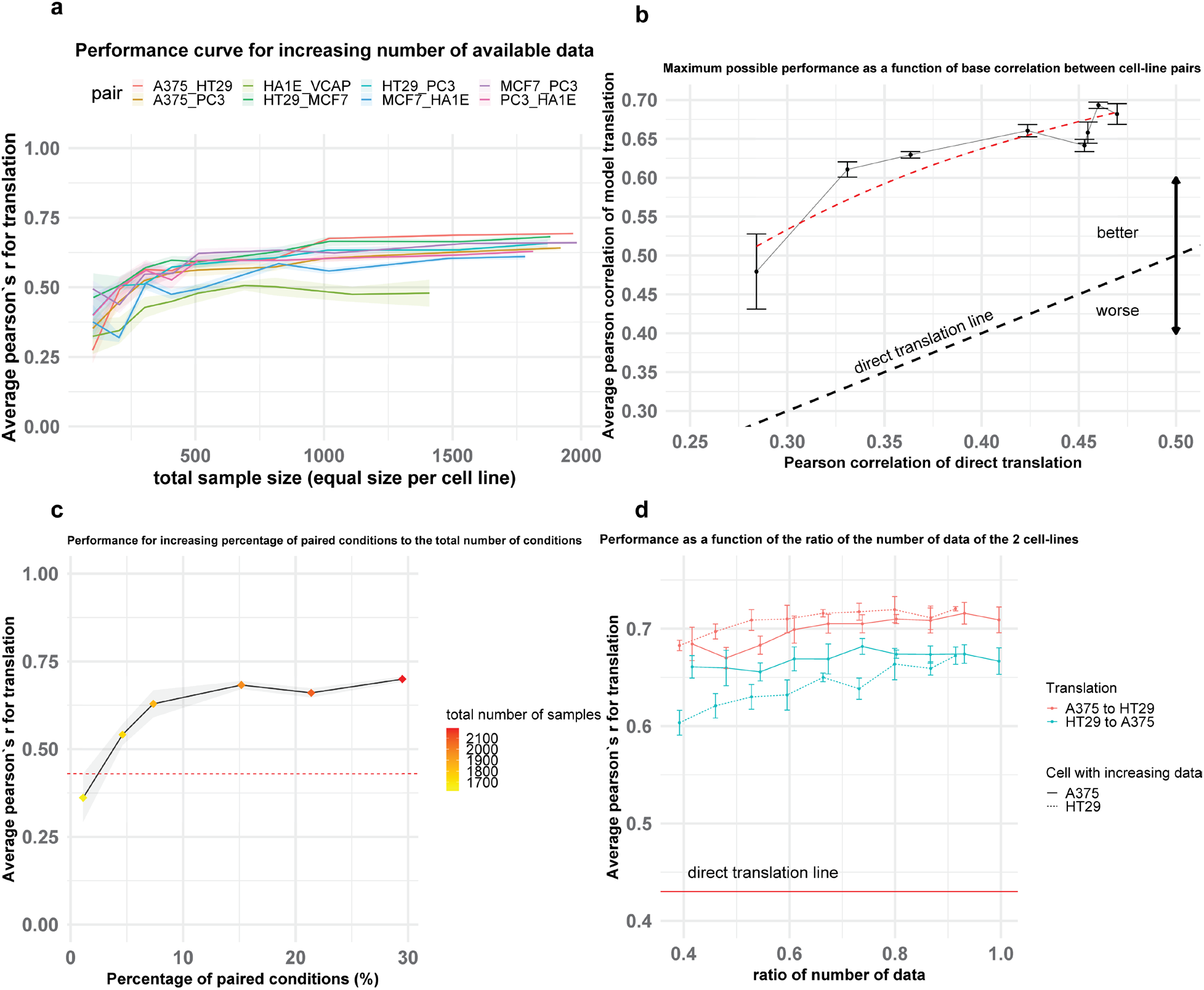
Analysis of framework’s performance and dependence on the data. **a)** Performance in the translation task of the CPA-based approach across different cell-line pairs and different sizes of training data. **b)** Model performance in translation as a function of the initial similarity of 2 cell lines. **c)** Model performance in translation for different percentages of paired conditions. **d)** Model performance in translation for low-to-medium cell-line imbalance in the conditions of the training samples.

### Evaluation of latent space embeddings

A global latent space is expected to have several properties to be suitable for translation. We evaluate the embeddings produced from our framework based on three criteria (Figure 3a-3c): i) different cell lines should not occupy different subspaces, so embeddings of pairs coming from the same cell line should not be more similar to each other than embeddings from random pairs, ii) pairs of embeddings coming from the same condition, regardless of cell line, should be similar, and iii) biological replicates should give similar embeddings, so pairs of embeddings from biological duplicates should be similar to each other. We evaluated these criteria using the cosine distance in latent space. No cell-line effect is observed in the global latent space, both for training and validation embeddings (Figure 3a, Supplementary Figure 8). Embeddings coming from the same condition are closer to each other than embeddings coming from random pairs (Figure 3b), while biological duplicates are even closer (Figure 3c), validating that indeed we have successfully constructed a stimuli-specific global latent space. Similar patterns can be observed in the global latent space when using the CPA-based approach (Figure 3d), but with a cell-line effect visible in the composed latent space, as expected with this method. We use Cohen’s d to quantify the difference between the distributions of cosine distances across all folds in 10-fold cross-validation (Figure 3e), proving that indeed there is a much higher cell-line effect in the latent space than the effect in the global latent space.

**Figure 3:**
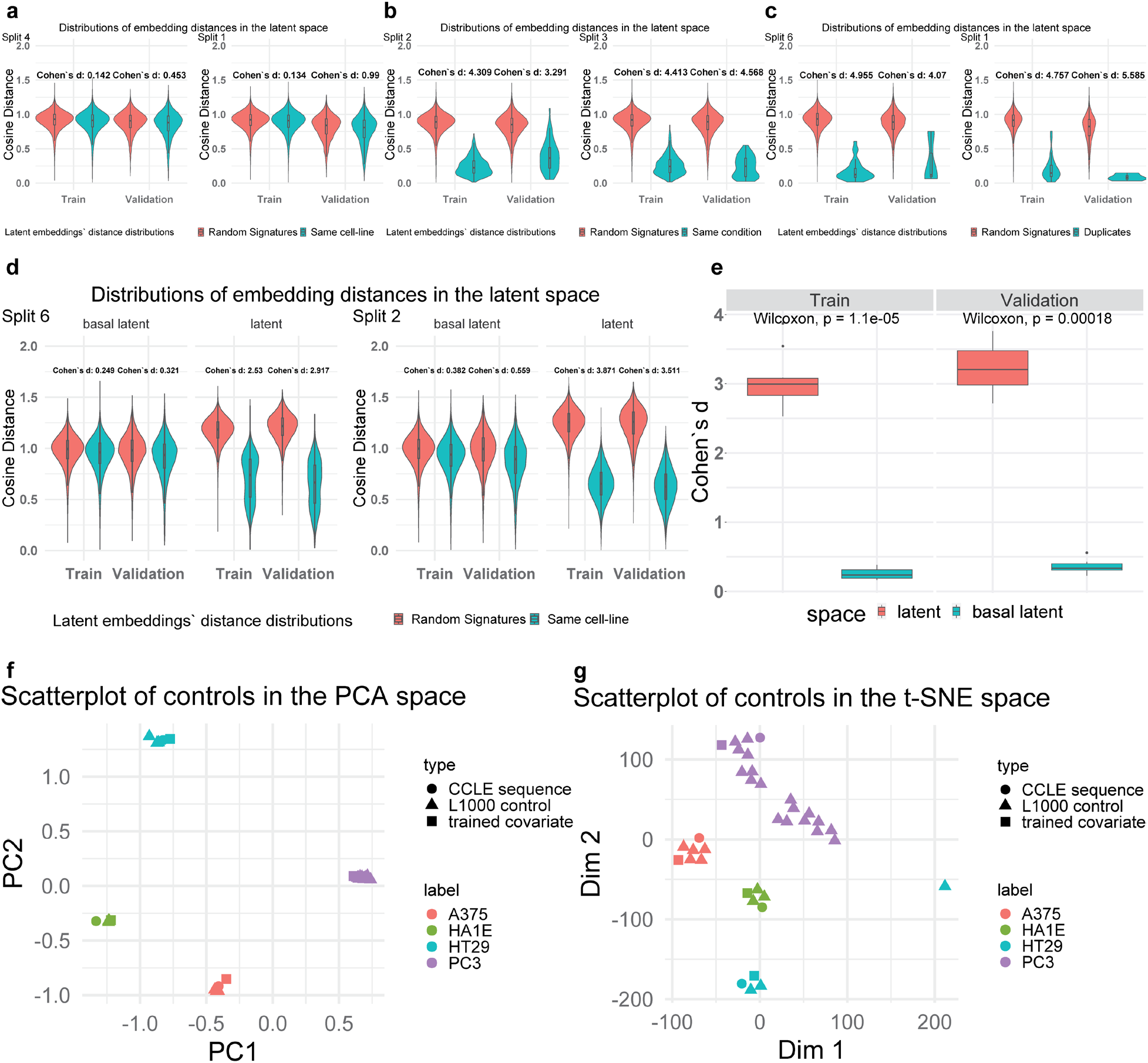
Properties of the latent space and model parameters interpretation. The two splits in 10-fold cross-validation present each time here are the ones where the maximum and minimum difference between the two distributions is observed. For every other split, the difference is between these two extreme cases. Additionally, panels a-c come from the first variation of the model with one global latent space while the rest come from the CPA-based approach **a)** Cosine distance between embeddings coming from random pairs of signatures and pairs coming from the same cell line. **b)** Cosine distance between embeddings coming from random pairs of signatures and pairs coming from the same condition tested on a different cell line **c)** Cosine distance between embeddings coming from random pairs of signatures and pairs being biological duplicates **d)** Distance between embeddings coming from random pairs of signatures and pairs coming from the same cell-line in the global and then the composed latent space in the CPA-based approach. **e)** Cohen’s d between distributions of cosine distances between random pairs of embeddings and embeddings coming from the same cell distribution. **f-g)** 2D-Visualization of L1000 control conditions, untreated cell lines from the CCLE dataset, and the trainable vectors of the CPA-based framework containing the cell line basal effect added to perturbations.

### Interpreting the biological information captured in the parameters

Deep learning models are often criticized for their lack of interpretability, so we investigate the biological information captured by some of the model’s parameters. Since only the cell-line effect is minimized in the global latent space of the CPA-like framework, the trainable covariate (covariates such as species, cell type etc.) vectors should only add a cell-specific effect. Intuitively, the global latent embeddings are expected to capture a “zero”/basal cell state corresponding to expression of untreated cells (controls), and thus the trained covariate which is added to that global representation should be similar to the composed latent space vectors which now captures the cell line effect. To investigate this we used control samples from the L1000 dataset not seen by the model during training, as well as samples coming from untreated cell lines from the Cancer Cell Line Encyclopedia^40^ (CCLE), using only the genes included in the L1000 landmark genes. Additionally, for this investigation two models were trained completely separately: the original benchmark model of A375/HT29 cell lines and another model using the PC3 prostate cancer cell line and the HA1E normal epithelial cell line. The latter pair was chosen because of high model performance (Figure 2a) and because these two cell lines are significantly different in terms of biology. Each trained covariate, even though the models were trained separately, is observed to be closer to its respective cell-line control signatures, both when using PCA for dimensionality reduction (Figure 3f), where clearly defined cell-line specific regions are observed, as well as when using t-SNE (Figure 3g). This demonstrates that some parts of the model are biologically interpretable and capture specific information.

### Identification of features that are important for translation and cell classification

The framework can be used to identify latent variables and genes that can be of biological importance. As a case study, we selected the model of the PC3 and HA1E cell lines with a classifier trained simultaneously to classify the cell lines from which the samples were derived (contradictory learning tasks). To identify the importance of genes according to the model for a variety of tasks with respect to their output, an integrated gradient-based approach^35^ was utilized (Methods) that attributes an importance score to each variable of interest. Since the same genes are used for both cell lines, it can be interesting to identify which are important for the model to translate a gene expression profile from one cell line to another cell line. Interestingly, the model attributes more importance to many genes other than the gene of interest when translating to across cell lines for the same condition (Figure 4a). In the case of the landmark genes, that phenomenon is slightly less prominent (Figure 4b). This is particularly interesting since one of the selected cell lines is cancerous and the other is non-cancerous, suggesting that the model may avert the fallacy of using the same gene as a proxy for its gene expression across disparate biological systems. Additionally, the model does not just attribute importance to genes that are highly expressed, based on Spearman’s correlation between the absolute importance scores and the absolute gene expression (Figure 4c).

**Figure 4:**
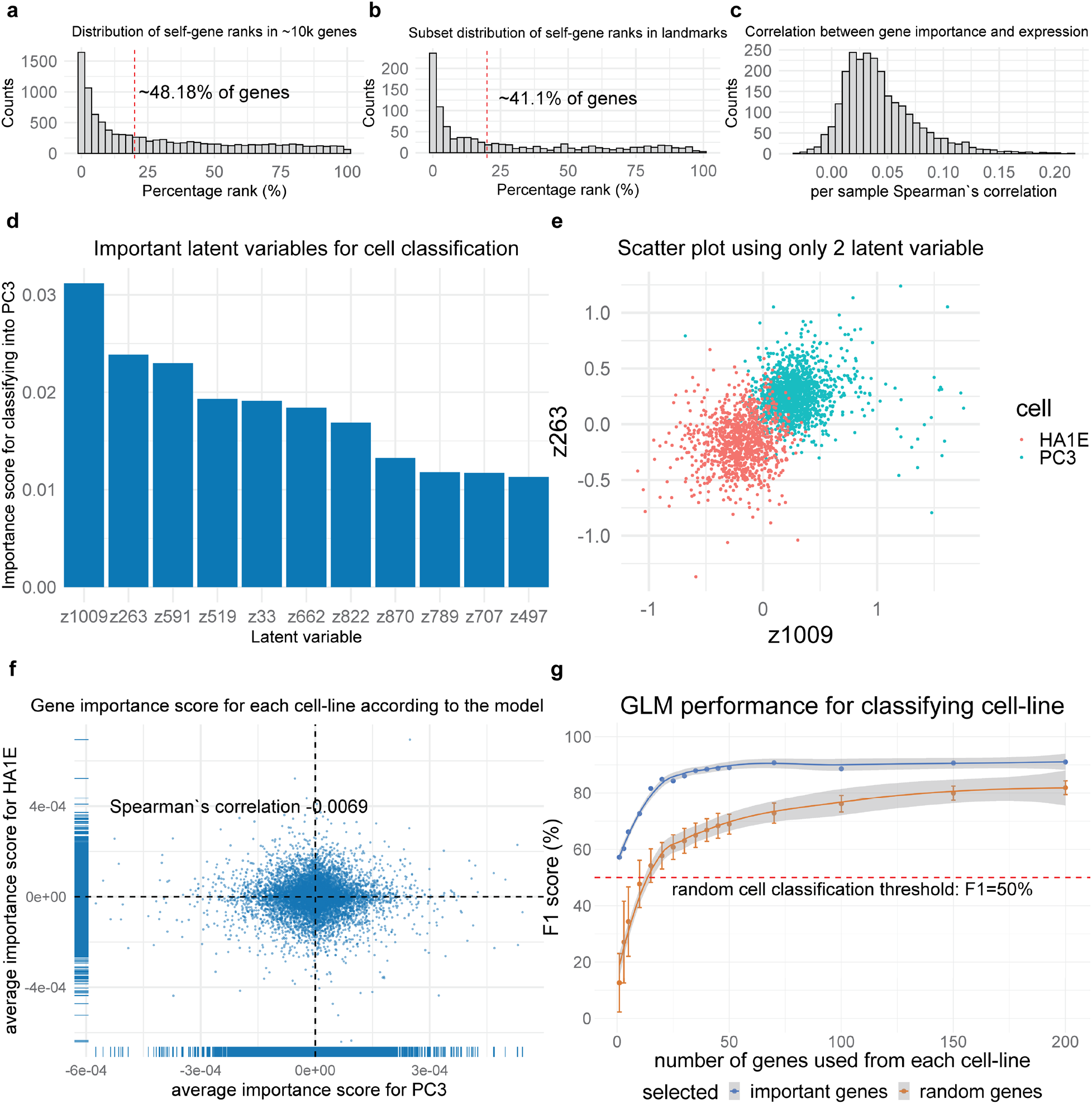
Feature importance investigation a-b) Distribution of percentage rank in terms of the importance of a gene to translate its expression to itself, using the 10,086 genes and the 978 landmark genes respectively in the L1000 dataset. **c)** Per sample Spearman’s correlation between absolute gene importance scores and absolute gene expression. **d)** Important latent variable to classify a sample as PC3 or HA1E in the global latent space, when a classifier is simultaneously trained. **e)** Separation of cell lines based on the top 2 most important latent variables according to the classifier. **f)** Average importance scores of genes from PC3 to control cell-specific latent variables versus the importance scores from HA1E, according to the individual encoders. **g)** Generalized Linear model classification performance by using increasingly more important genes.

The simultaneously trained classifier can also be used to identify subsets of latent variables in the global latent space that are important for classifying samples by cell type. Although the cell line effect is partially filtered and embeddings coming from the same condition are globally close to each other (Supplementary Figure 4), there are still 11 latent variables that allow the classification of cell line (Figure 4d) using a k-means-based approach (see Methods). These latent variables can separate the samples based on cell line (Figure 4e), even though globally the cell line-specific effect in the latent space is still filtered out. Genes considered important by the encoders to control these latent variables should be either cell line-specific genes or a subset of genes that can easily distinguish between cell lines. The importance scores of the genes for each cell line-specific encoder do not correlate at all and are different between the two cell lines (Figure 4f). It is possible to even train a very simple generalized linear model to classify cell lines based on gene expression, only using a subset of these important genes, achieving high performance with only few genes from each cell line (Figure 4g).

### Performance in inter-species translation for lung fibrosis

Animal models don’t recapitulate human biology perfectly, so computational modeling can be used to improve the translation between humans and animal models. We evaluate the ability of the framework to perform inter-species translation. We utilize the raw gene counts coming from single-cell RNA-sequencing of a mouse^36^ and human^37^ lung fibrosis dataset. Similar to the original CPA study^32^, the decoders predict a mean and a variance for every gene, derived from a negative binomial distribution. Furthermore, both a trainable species vector and another trainable cell type vector are added to the global space, in attempt to minimize both species and cell type effects. We evaluate the performance in the reconstruction of gene expression profiles and the ability to translate between mouse and human under 10-fold cross-validation in terms of R^2^ of the predicted per gene means and variances, where we would expect to observe a similar distribution in a successful translation, and thus mean and variance. Our framework outperforms the other approaches in terms of R^2^ of the means both in reconstruction and translation (Figure 5a). When predicting the within-gene variance, there is not always a statistically significant improvement, as all approaches have generally low performance (Figure 5b), which suggests that the models fail to capture variation in gene expression. We do not find any significant difference in performance between using all genes or just homologs (Figure 5a-5b). It is worth noting that based only on the human lung fibrosis dataset, three of the top ten genes contributing to the top principal components do not have homologs in mice (Supplementary Figure 11), meaning that irrespective of performance, a method considering only homologs would exclude important genes for lung fibrosis.

**Figure 5:**
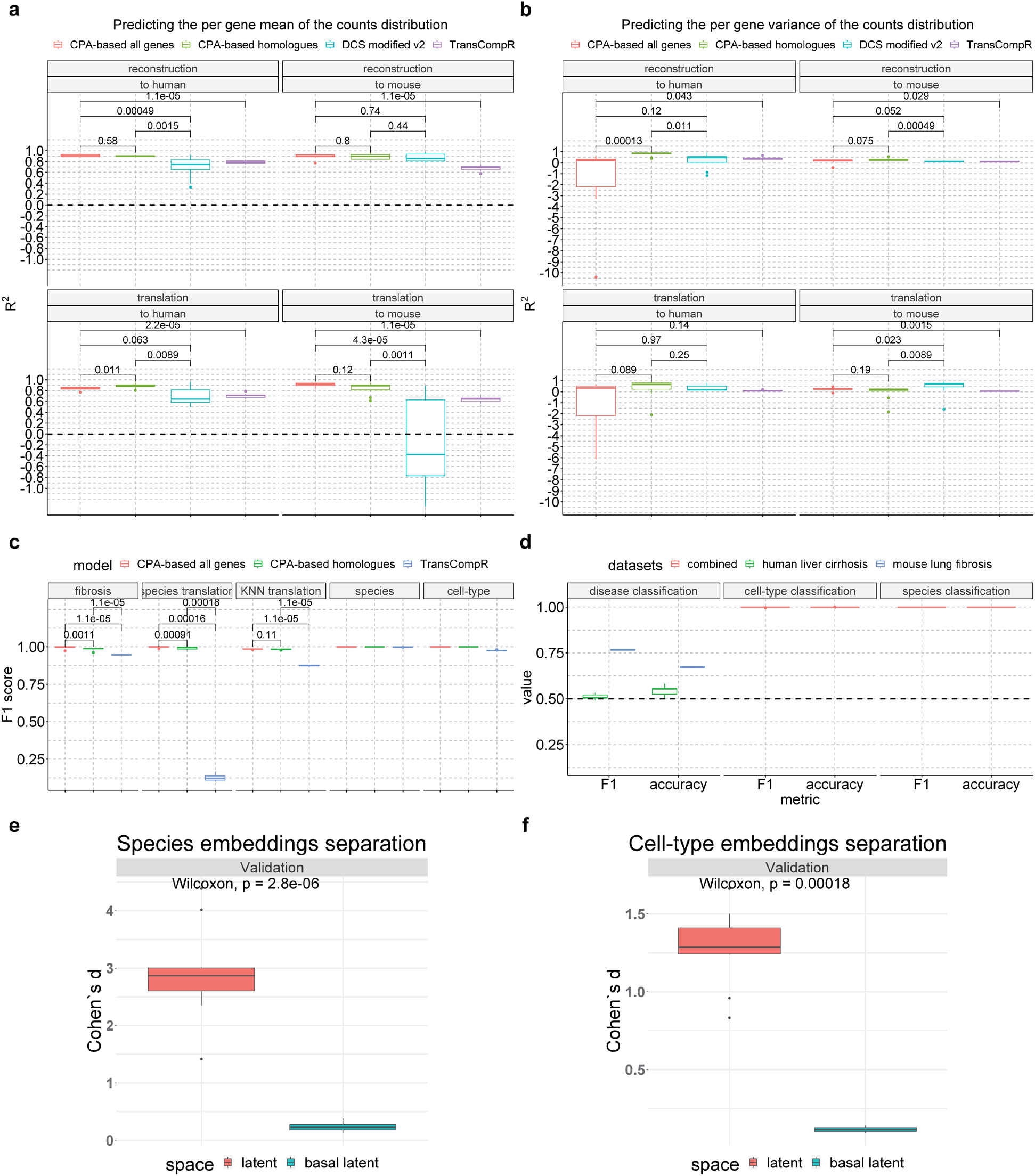
Evaluation of the framework in inter-species translation in fibrosis a-b) Performance (R^2^) in predicting the per gene mean and variance of single-cell RNA-sequencing data for the tasks of reconstruction and species translation in the human-mouse lung fibrosis datasets. The comparison is done between the CPA-based framework using all genes or homologs and TransCompR **c)** Classification performance comparison in different tasks. **d)** Classification performance of the CPA-based framework using all genes in external disease datasets. **e)** Embeddings separation based on species in the global latent space versus the composed latent space. The effect size d is calculated as Cohen’s d. **f)** Embeddings separation based on the cell type in the global latent space versus the composed latent space. The effect size d is calculated as Cohen’s d.

We also evaluate the ability of each approach to classify fibrosis, species, and cell type and to classify correctly a signature as a different species when that is translated in the composed latent space, by adding a different species effect. In our framework, utilization of all genes outperforms the homolog genes approaches in predicting fibrosis and species-translation, though the performance of all approaches is high (Figure 5c). Similar to what was observed for the L1000 dataset, species and cell type are perfectly predicted in our framework. Additionally, both the species and cell-type effects are filtered (Figure 5e-5f, Supplementary Figures 9-10) in the global latent space compared to the composed latent space, meaning the model succeeds in removing the cell type and species effect in the global latent space and then retrieving it again in the composed latent space.

### Generalization in other disease datasets

Models that are trained on a specific data set can often perform worse on external test sets, and it is, therefore, useful to investigate to which extent the model can predict disease, species, and cell types in other datasets, as well as different tissue and disease datasets. For this, we use an independent dataset on mouse lung fibrosis^41^ and a dataset on human liver cirrhosis^42^. In the mouse dataset, even though different genes were measured than those in our model, the performance is still decent in disease classification (Figure 5d). For the human dataset, which is an extreme case of fibrosis in a different organ, the model has markedly lower performance although better than chance (Figure 5d). Interestingly, in both cases, the model can still perfectly identify cell types and species (Figure 5d), once again displaying the model’s ability to capture the general characteristics of the system.

### An inter-species model from serology data for predicting protection against HIV

As a final case study, we developed a model for cross-species translation of serology data, where there are no 1-1 mapping of features, to predict vaccine-induced protection from HIV in humans. Previous failed HIV vaccine trials have suggested that neutralizing antibody titers, the primary outcome for most vaccine trials, do not consistently correlated with vaccine efficacy. Moreover, recent research suggests that deeper characterization of the antibody response, including antibody subtype prevalence and Fc-receptor binding affinity, may be necessary to predict the quality of the vaccine response. Notably, the primary challenge in comparing pre-clinical animal models and human clinical trial data in this context is that antibodies and Fc-receptors with similar names across species can be structurally and functionally distinct and orthologous features might not exist. Our ANN approach has the potential to advance our understanding of which preclinical features might best predict efficacy of a HIV vaccine. Here we utilize serology data from non-human primate (NHP) and human datasets ^38,39^ following vaccination against SHIV and HIV, respectively. In this case, we added a non-linear species effect using two small intermediate fully-connected neural networks with non-linear activation functions (Figure 6a). In line with other models constructed using this framework, the model was trained so that protected individuals are close in the global latent space regardless of species. While two separate classifiers try to predict vaccination status and protection in the global space (Figure 6a), a third classifier predicts species in the composed latent space. For the human serology features, the model has high performance when reconstructing each feature (Figure 6b, r = 0.89±0.01). Ιn NHPs, while some features are not predicted well and there is a big variation in performance between folds, the overall performance is still good (Figure 6c, r = 0.71±0.04). Finally, the performance across all classification tasks is exceptionally high (Figure 6d) including 100% accuracy in species classification and translation, which is evaluated by how well the species classifier predicts species label when translating a signature to another species in the latent space.

**Figure 6:**
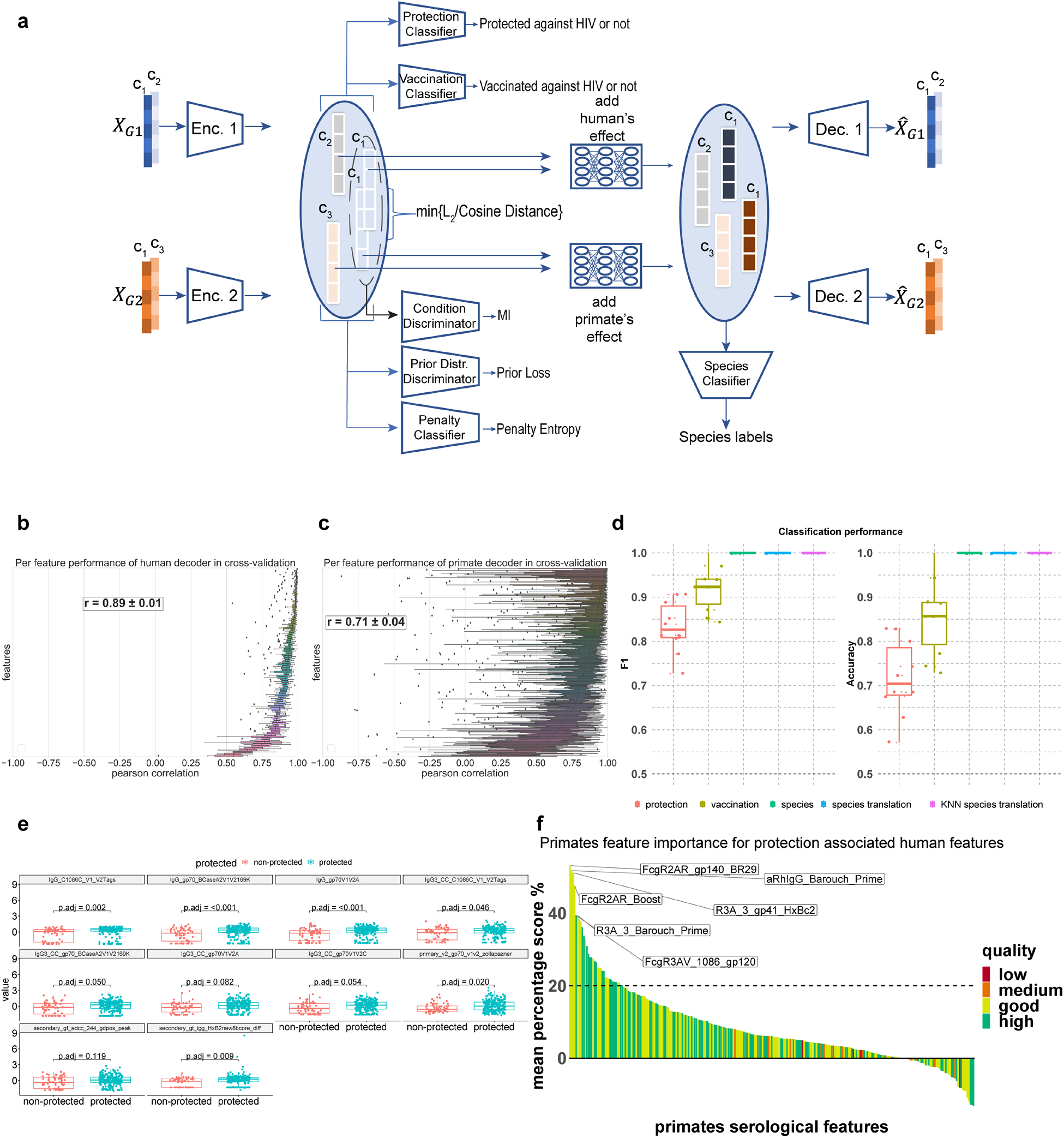
Inter-species translation of serology data. **a)** Framework architecture for inter-species translation in the serology dataset. Instead of adding the species effect with a trained vector, it is adding completely non-linear with 2 small intermediate neural networks. **b)** Per feature Pearson correlation across in 10-fold cross-validation for human’s features. **c)** Per feature Pearson correlation across in 10-fold cross-validation for non-human primates’ features. **d)** Classifiers’ performance in various tasks. **e)** Univariate differences between protected and non-protected individuals, for human serological features related to viral protection, as found from the integrated gradient and LRT approach. **f)** Non-human primate features predictive of human protection, by using importance score for translating them into human features. Their quality is assigned based on the average reconstruction Pearson’s r of these features in 10-fold cross-validation: i) low quality = 0<r<0.25, ii) medium quality = 0.25≤r<0.5, iii) good quality = 0.5≤r<0.75, iv) high quality = r≥0.75

Using the model, we aimed to identify features from both species that are predictive of human protection. For this, we performed the integrated gradient approach in parallel to likelihood ratio tests (LRT) on each latent variable (see methods). Latent variables are denoted as important in predicting human protection only if there is an agreement between the likelihood ratio test results and the integrated gradients (Supplementary Figure 17). The human features identified indeed have a statistically significant difference between protected and non-protected individuals (Figure 6e). Finally, we identified NHP features that have a high gradient score when translating to human signatures, meaning these NHP features are predictive of human features linked with viral protection (Figure 6f). These features are not necessarily associated with NHP protection (Supplementary Figure 18) but they could be predictive of human protection. Notably, while the top human features identified are generally related to IgG titers, the top NHP features are mostly related to the Fcg-2A and Fcg-3A receptors. Further analysis of feature importance could potentially identify a set of serological biomarkers in NHPs that is highly predictive of human HIV vaccine efficacy.

## Discussion

Here we develop AutoTransOP, an ANN framework that facilitates the translation of omics profiles between different biological systems. The framework builds upon ideas of the CPA approach^32^ and other species and cellular translation methods^13,15,18,31^, together with ideas from language translation models^33^. The explicit goal is to align omics signatures between systems, rather than identifying what information inherent in the signature of one system is most germane for understanding phenotype characteristics in the other, which has been the objective in many previous studies^16–19^. The framework performs as well as (or even better than) other state-of-the-art translation techniques, when using homolog features between systems, and performs similarly also without a 1-1 mapping between features. Notably, the framework constructs a truly global latent space with stimuli-specific regions, for which classifiers can be jointly trained to make predictions for various tasks such as the diagnosis of diseases.

Most current approaches to translating between systems require homolog features and utilize linear transformations to facilitate translation^13–18^, and are thus restricted to represent linear inter-species relationships. Also, the non-linear ANN-based approach DeepCellState^31^ requires homology of the molecular features used to describe the biological systems. In contrast, our framework can represent non-linear relationships between different biological systems, without requiring any kind of homology, and achieves high performance using only a small percentage of paired conditions. This enabled us to train a translation model on serology datasets for which a 1-1 mapping of the features between the two biological models did not exist. Through interpretation of this model, relationships between very different molecular profiles that correlate with specific phenotypes can be identified, e.g. protection against infection.

Interpretability of deep learning models in biology remains a challenge. These models have been criticized for providing a poor understanding of which biological relationships they capture ^43,44^. On this front, we demonstrate in our framework how integrated gradient approaches^35^ can be used to estimate the importance of features used by different parts of the framework for various tasks, enabling some biological interpretation of the model. Based on this, we could propose serological features predictive of human protection against HIV, including non-human primate-specific features that can be observed in preclinical stages of vaccine development. Finally, elements of the framework can be used to interpret and successfully retrieve the effects of species or cell types, filtered from the global latent space. This can explain the ability of the framework to predict cell types and species with high performance also in independent disease datasets, derived from different organs/tissues. However, there are still limitations in the generalization of in the models to external datasets. In particular, the performance on such datasets drops significantly as samples from different pathologies and tissues are considered. Even within the same disease, the inclusion of different features can lead to reduced performance in predicting disease diagnosis.

Despite our framework being trained successfully on datasets with relatively small sample sizes, the model still contains many parameters, especially when using a larger number of features, which inevitably leads to overfitting. Some of these shortcomings could likely be alleviated by applying our framework to larger datasets, such as ARCHS4^10^, which contains hundreds of thousands of publicly available RNA-sequencing data from humans and mice. Training with more data and more diverse unique conditions may enable higher generalization and higher granularity in modeling different biological covariates. Additionally, with the advent of Natural Language Processing (NLP) models^45^ and attention-based models^46^, our encoder modules could potentially be modified with NLP-like representations. Recently, Geneformer^47^, an attention-based model, was pre-trained on a corpus of 30 million single-cell transcriptomic profiles and was proven to be context-aware of the system it encodes. Although it still requires some level of homology, it paves the way to utilize NLP approaches for transfer learning in biology, and ultimately translation.

The flexibility of our framework allows the modeling of many different biological systems. This could lead to the computational optimization of biological systems and assays aiming to model human pathology. Using our framework, we can both explore potential transcriptional modifications to design better disease models and identify features predictive of human biology without requiring homology between systems, ultimately reducing resources spent during experimental modeling and potentially expediting the translation of in-vitro and pre-clinical findings to human therapeutic advancement.

## Methods

### Pre-processing of *in-vitro* transcriptomics benchmark dataset

The L1000 CMap resource^12^ contains bulk gene expression data from drug perturbations across different cell lines and provides a benchmark dataset with diverse conditions and a large sample size (for a total of 720,216 samples of drug perturbations of varying quality). Additionally, several equivalent perturbations across different biological systems are available (406 *paired conditions* for the case of A375 and HT29 cell lines after filtering and pre-processing, explained below) to evaluate the performance in translating omics profiles. We selected high-quality drug perturbations from the latest version of the L1000 dataset (accessed via clue.io). The level 5 z-score transformed and pre-processed differential gene expression data of 978 landmark genes, measured with the L1000 assay, and additionally, 9,196 computationally inferred genes in the CMap resource that were marked as well-inferred, were considered in the subsequent analysis. We consider perturbations as high-quality if they consist of signatures with more than three replicates, where at least half of them passed the standard quality control protocols in the assay, as provided in the dataset, and were not identified as statistical outliers (as considered by the L1000). Additionally, where multiple-signature perturbagens, i.e. technical duplicates, only the signature with the highest transcriptional activity score (TAS) across these technical duplicates was retained in the dataset, these signatures are labeled ‘exemplars’ by the CLUE platform and are specifically designated for further analysis by the platform^48^. The TAS metric is provided along with the L1000 dataset and quantifies signal strength and reproducibility. Finally, the ability to distinguish between random signatures and true biological duplicates, meaning the same perturbagen tested on the same cell line for the same duration and dosage, was evaluated for different parts of the dataset, split using varying TAS thresholds (Supplementary Figure 19) and samples with a TAS≥0.3 were retained. After filtering and 13,699 samples remained, with 1107 conditions available in total for the HT29 cell line and 1213 for the A375 cell line. In the case of control signatures, we followed the same procedure but without filtering based on TAS.

### Pre-processing single-cell RNA sequencing interspecies datasets

For the human and mouse single-cell RNA-sequencing datasets, we first re-annotated manually each annotated cell into one of the four classes: i) immune cells, ii) mesenchymal cells, iii) epithelial cells, iv) endothelial cells, and iv) stem cells. These high-level labels were later used to remove cell effect from the global latent space and were also used in the subsequent cell-type classification. Finally, while the raw gene counts are used for reconstruction from the decoder in the loss function, the encoders take as input the log-transformed counts (*x_i_*_*input*_= log_10_(*count* + 1)), which acts as an activation function in the first layer of the encoder.

### Pre-processing of the serology datasets

For all serology data, we aimed to construct a model using only antibody and receptor measurements. The human data were retrieved from Chung, Kumar, Arnold et al.^39^ upon request, the avidity molecular features were dropped and the data were z-scored per feature. The non-human primates’ data were retrieved from Barouch, Tomaka, Wegmann, et al.^38^ upon request, the samples taken in week 28 were used, and antibody-dependent cellular function features and mass spectrometry data were dropped. The data were log-transformed (*x* = log_10_(*MFI* + 1)), the median per feature from controls is subtracted from each feature to standardize the data. Finally, the data are z-scored per feature.

### The general framework and the training procedure

In this implementation, the framework always models pairs of systems for translation, species, or cell lines. Each is modeled with separate encoders and decoders for each of the species or cell lines in the pair attempting translation, while inside a latent module, the global latent space is shaped (Figure 1a). Both the encoders and the decoders are multi-layered neural networks, with each layer consisting of, sequentially: a fully-connected layer, a batch normalization layer^49^, an ELU activation function^50^, and a dropout layer^51^. The final output layer of the encoder and the decoder consists of only one fully connected layer without a trainable bias term.

For the construction of the global latent space several metrics are optimized: the distance (***L*_*distance*_**) between embeddings of profiles coming from different systems undergoing the same perturbation is minimize and their cosine similarity (***L*_*cosine*_**) and mutual information (***L*_*MI*_**, see details below) is maximized; and the divergence of the distribution of the latent variables from a random uniform distribution is minimized (***L*_*prior*_**). Both cosine similarity and Euclidean distance losses were added to enforce the strongest possible filtering of species and cell type effect, while the cosine similarity also enforces normalization of the latent embeddings. This is achieved using two different ANN discriminators, as previously proposed in the MINE^52^, Deep InfoMax^53^ and InfoGraph^54^ studies, where the Jensen-Shannon Mutual Information between embeddings coming from the same perturbation is estimated and the extra prior loss is calculated and added in the final loss, according to the following equations with the implementation taken from the deepSNEM model^55^.

- 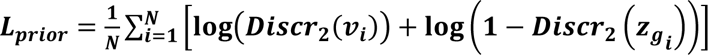 where *v*_*i*_ is a randomly sampled embedding from a prior random uniform distribution ranging from 0 to 1 and *z*_*g_i_*_ is a global latent space embedding. N is the number of samples in a batch during training.
- 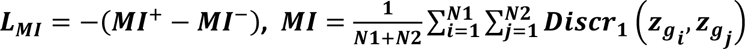 where MI^+^ and MI^-^ are respectively the mutual information between pairs derived from the same conditions and pairs coming from different conditions, averaged for every possible pair in a batch during training. *z*_*g_i_*_, *z*_*g_j_*_ are global latent space embeddings, whose mutual information is estimated by the Discr_1_ discriminator.

Discr_1_ is the discriminator estimating the mutual information between two embeddings from the global latent space. It takes as input two global latent space embeddings and passes them through the same 3 fully-connected layers, each of them followed by a ReLU activation function^50^ and one fully-connected skip connection. Then the product of the result of this non-linear transformation of the two embeddings is used to approximate a lower bound of their Mutual Information, as proposed originally in MINE^52^ and Deep InfoMax^53^. Discr_2_ is the second discriminator which takes as input an embedding vector and calculates the probability a point in this embedding space is sampled from a specific distribution. This way *L*_*prior*_ forces each feature of the learned embeddings to be sampled from a distribution close to the random uniform distribution ranging from 0 to 1. It has three similar fully-connected layers and the final scalar output is passed through a sigmoid activation function^50^. These regularization loss terms (***L*_*distance*_, *L*_*cosine*_, *L*_*MI*_**) are calculated and averaged across every pair of global embeddings (*z*_*g_i_*_, *z*_*g_j_*_) that are coming from the same condition. The ***L*_*prior*_** is calculated for every sample in the dataset, meaning every global latent embedding and averaged across samples. For the case of the L1000 dataset, we consider similar perturbations those that are coming from experiments of the same drug, tested on the same cell line, with the same dose and time duration. For the lung fibrosis dataset, similar profiles are considered those coming from samples that have the same diagnosis (fibrosis or not). For the serology datasets, we train the framework so that embeddings coming from protected individuals against HIV are close to each other regardless of species (and even vaccination status)

The basic task of this autoencoder framework is reconstruction, which is achieved by minimizing some kind of reconstruction loss (***L*_*recon*_**). In the case of z-scored profiles from bulk data, this is done by minimizing the mean sum of squared errors between the input of the encoders and the output of the decoders. The sum of squares error is averaged across samples. For only the case of single-cell RNA-sequencing data, based on the implementation proposed in the CPA manuscript^32^ (found here https://github.com/facebookresearch/CPA), the negative binomial negative log-likelihood is used to optimize the reconstruction, by assuming that the data are derived from a negative-binomial distribution characterized by a mean and variance that are both predicted. The negative binomial negative log-likelihood loss is calculated for every sample and the average across all samples in the batch is minimized.

Classifiers are used for different classification tasks. These consist of multiple fully connected layers and a final SoftMax activation function before the output. The average entropy loss across samples for every classification task in the latent space is minimized: *entropy* = 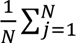 *CrossEntropy*(*Classifier_i_*(*z_j_*), *label_j_*), where *entropy* is the average cross entropy between every j^th^ prediction of a classifier taking a latent vector as input and the true label for that sample.

L2-regularization of the weights and bias of the encoders (***L*2_*encoder*,*i*_**), decoders (***L*2_*decoder*,*i*_**), and classifiers (***L*2_*classifier,i*_**) is also enforced by minimizing the sum of squares for the aforementioned trainable parameters.

Taken together, for the basic variation the following loss function is optimized

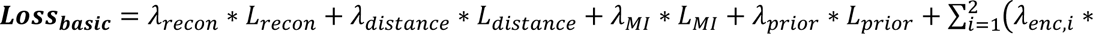 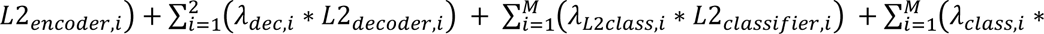 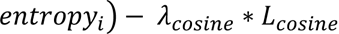, where M is the number of classifiers and thus individual classification tasks, and the rest of the terms, together with how they are calculated, have already been described in the previous paragraphs of this section. For values for each of the λ used in the loss function see Supplementary Table 5.

### Variation of the global latent space with a simultaneously and competitively trained classifier

For the variation of the global latent space with a simultaneously and competitively trained classifier the aim is to embed some species or cell line information in some of the latent variables. A simple classifier for correctly predicting the cell line label is trained simultaneously on the global latent space with the rest of the framework and an entropy loss is added to the original description of the framework. The construction of a global latent space and the training of the classifier are competing tasks, where the framework is trained to achieve a stable trade-off.

### CPA-based variation of the framework

For the CPA-based framework, the global latent space is expanded by augmenting the loss function with some additional terms.

An adverse classifier of species and cell types is added. As described in the original CPA manuscript^32^, during training we iterate between training the classifier (updating only its parameters) on the global latent space, and training the rest of the framework with the addition of a penalty (***entropy_adverse_***) if the classifier correctly classifies species and cell types. To improve the robustness of the discriminator it is initially pre-trained only with encoders and discriminators, without other classifiers and the decoders, so that it can already distinguish cell types and species in the global space.

Furthermore, species and cell type effects are added to the latent space via trainable vectors. In the newly composed latent space, from which the decoders are sampling embeddings, classifiers are jointly trained to correctly classify cell types, and species (or even disease diagnosis). Additionally, the trainable vectors are regularized by the L2 norm (***L*2_*trained_effect_*_**). All the above can be summarized in this new loss function:

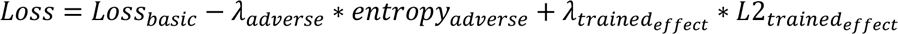

### Framework for the serology datasets

In the serology dataset we utilize the CPA-based framework but instead of adding the species effect with trainable vectors, small artificial neural networks are utilized consisting of two fully-connected layers. L2-regularization terms for these small ANNs are added to the training loss function.

Additionally, it is aimed to later identify features predictive of protection or vaccination status regardless of species. For this purpose, we train two classifiers predicting vaccination and protection status in the global latent space. We care more about protection and thus, as described previously, we aim to create similar embeddings and minimize their distance in the global latent space just by looking at protection status.

### Framework’s basic hyperparameters

Here we present the basic parameters used to train the model. No thorough hyperparameter tuning was performed, and values were selected based on empirical values and tuned so that there is convergence in the training loss and the training reconstruction performance (Pearson’s r and/or R^2^). Additionally, these values were also tuned so that the performance in training (not validation) is sufficiently high, meaning that the model is at least able to fit the given data. This empirical tuning was done only based on the 1^st^ training set in 10-fold cross-validation.

The latent space dimension was chosen to be as small as possible until the model’s performance dropped in both training and validation of only the 1^st^ fold. Based on this latent dimension and the original input dimension of the data the sizes of hidden layers of the encoders were chosen to be in-between, gradually reducing the input dimension to that of the latent space. The actual size and number were constrained by practical memory limits. The decoders had the same number and sizes of hidden layers as those of the encoders, but now they increase the size of the embeddings from the latent dimension to the original input dimension.

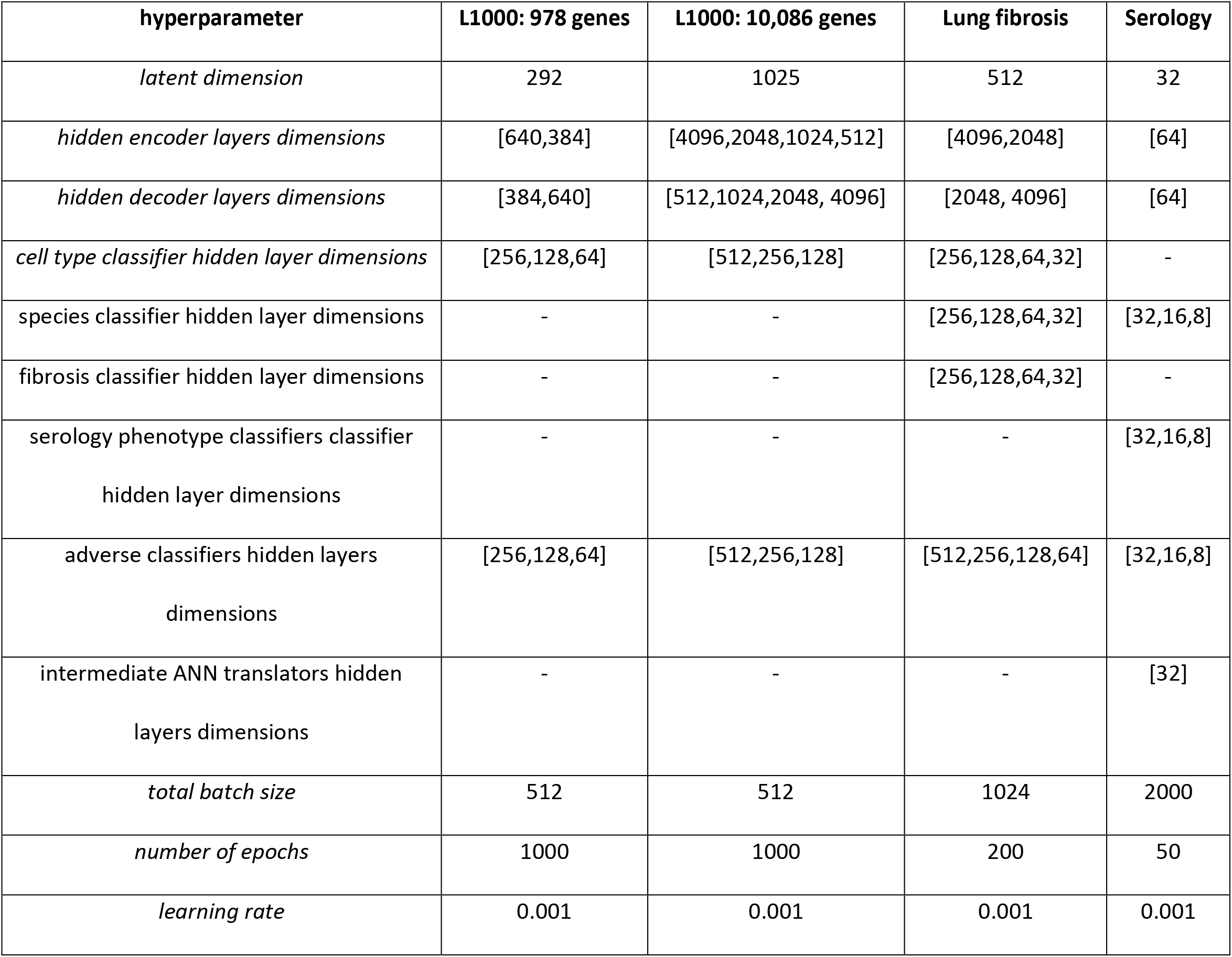

### Evaluation procedure and metrics

The model performance was evaluated using 10-fold cross-validation. One fold of the data was hidden during training and used to evaluate performance in unseen data, and 90% of the data from each system (species or cell line in the case of L1000) were. For the L1000 dataset, for evaluating the translation of the whole omics profile, we made sure that for the case of paired conditions, the perturbation in both cell lines was hidden during training.

The classification tasks were evaluated by total accuracy and F1-score (or micro F1 for multiple categories):

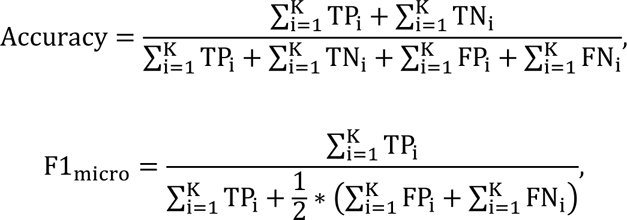

where K is the total number of classes in multi-class classification, TP and FP symbolize true and false positives, and TN and FN symbolize true and false negatives. For the case of multiple classes, we define as positives the samples belonging to that specific class while everything else is a negative sample. Using this definition of positives and negatives we further calculate the TP, FP, TN and FN per class. In the case of cell-type classification in lung fibrosis K=5.

For the cell line classification in L1000, species classification both in lung fibrosis and the serology datasets, and vaccination and protection status in the serology dataset we use the F1 score and accuracy for binary classification 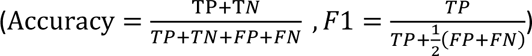.

To evaluate the validity of the predictions (*ŷ*) of whole signatures in translation and reconstruction, compared to the ground truth (*y*), we utilized:

i. the global Pearson’s correlation 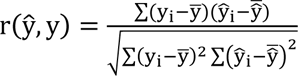, where ŷ and y are flattened and the i^th^ element is the i^th^ point in these flattened vectors.
ii. the average per sample Spearman’s correlation 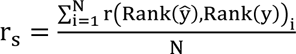, where N is the number of samples and Rank() means ranking the gene based on their differential gene expression and using these ranks to calculate Spearman’s correlation.
iii. the average per sample sign accuracy = 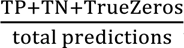, where TP signifies the genes that total predictions have a positive sign regulation both in the actual data and predictions, TN signifies the genes in the sample that have a negative sign regulation both in the actual data and predictions, and TrueZeros are the genes that have an absolute expression ≤10^-6^ both in the actual data and predictions (a small tolerance rather than strictly zero was chosen for numerical reasons).

For the single-cell RNA-sequencing data where we predict the per gene mean and variance we calculate the coefficient of determination (R^2^) per gene mean and variance, similar to the original CPA manuscript^32^. In general, R^2^ is calculated as: 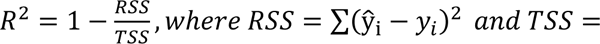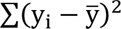

### Separation of latent space embeddings

To evaluate the similarity of embeddings for different signatures, and whether there is separation based on cell, species, or conditions in the latent space, we utilize cosine distance, ranging from 0 (the same) to 2 (completely) different: 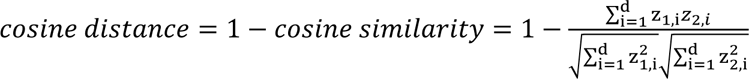, where z_1_ a and z_2_ are two latent space vectors to be compared and d is the total number of elements in the vector, i.e. the latent dimension.

To estimate if there is a cell, species, or condition effect, and compare it between the composed and global latent space we utilize Cohen’s d to estimate the effect size between the distributions of cosine distances, derived from random pairs of embeddings and pairs coming from the same cell, species, or condition. The effect size is thus calculated using the mean and standard deviations of two cosine distance (cos) distributions as: 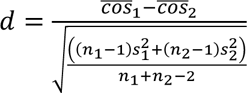, where n_1_, n_2_ is the number of samples of each ofthe two distance distributions, 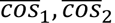 are the means of the cosine distance distributions and s_1_,s_2_ are the standard deviations of the cosine distance distributions. A Cohen’s d around 0.8 is a large effect size (around 2 is considered a huge effect size) while around 0.5 is a medium effect size, and around 0.2 and below is considered small or very small^56,57^.

### Feature importance using integrated gradients

To estimate the importance of features we utilize integrated gradients^35^ from the Captum library^58^.

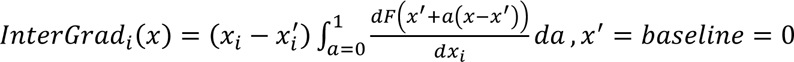

The importance scores are calculated based on the gradient with respect to the input of the model, and thus the higher the absolute integrated gradient the higher the importance of that input feature to control the output. A negative score means the variable has a negative effect pushing the prediction to the other class, while a positive score has a positive effect.

For example, if we want to identify important latent variables to classify a sample as one coming from a particular cell line we calculate the integrated gradient of every latent variable to make the classification and take the average across all samples. Similarly, if for example, we are aiming to calculate the importance of genes to control latent variables in the global latent space we can calculate the integrated gradient score of every gene for every variable in every sample, and then take the average across samples.

### K-means-based separation of important latent variables

Latent variables can be separated into important and unimportant ones using k-means, inspired by an approach that was used to identify important connections between latent components and genes in microbial organisms by using the weights derived from independent component analysis ^59,60^. We assume that only 2 main clusters of latent variables exist, one containing important variables and one containing unimportant ones. On this front, the latent variables are clustered based on their absolute gradient scores into 3 clusters, where 3^rd^ cluster is assumed to be a very small cluster of outliers. The midpoint between the variable with the highest score in the unimportant cluster and the variable with the lowest score in the important cluster is used as a threshold to distinguish between significantly important and unimportant latent variables. As a sanity check the important variables are also compared with the top-ranked variables based on their score.

### Likelihood ratio tests for the identification of important latent variables

To identify which latent embeddings correlate with viral protection after accounting for vaccination status and species, a likelihood ratio test (LRT) was performed on each individual latent variable. Here, the likelihood (L) of the alternative model (*H_A_*): *latent variable_i_ embeddings ∼ prtection + vaccination + species* was compared to the likelihood of the nested model, or null hypothesis, (*H*_0_): *latent variable_i_ embeddings ∼ vaccination + species* in 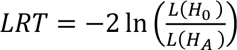. We rejected *H*_0_ for *latent variable*_i_* when the FDR-adjusted p-value of the chi-square test was less than 0.05, concluding that the model including protection has a statistically significant better fit than the model without protection. In the volcano plots, the − log(pvalue*) is plotted against the t-value for the protection term in the alternative model. This method assumes that the relationship between the latent variable embeddings and protection is linear. R package *lmtest*^61^ (version 0.9.40) was used to perform these statistical tests. Finally, the intersection of these latent variables with significant latent variables (average percentage importance score across folds ≥10%), based on their gradient score from the trained protection classifier, is used for the final identification of robust latent variables associated with viral protection. We keep latent variables that the sign of correlation with protection agrees in both approaches.

### Identification of protection-associated serological features

The importance of the serological features is calculated as previously described with the integrated gradient score of every feature for every latent variable that was identified to be statistically significant for predicting viral protection of humans, averaged across samples coming from the respective species. Serological features with high scores (and at least ≥20%) can control latent variables in the global latent space associated with human viral protection, and thus they are predictive of human protection. For human features, we also validate that the univariate differences between protected and unprotected individuals are indeed significant, by using a non-parametric Wilcoxon test, with Bonferroni correction for multiple hypothesis testing.

Finally, we calculate the integrated gradient score for translating each non-human primate serological profile to a human profile. The non-human features with high scores the human ones associated with protection, can be considered serological non-human features predictive of human viral protection.

### Inference of transcription factor activity

To infer the transcription factor activity, we utilized the VIPER algorithm^62^ together with the Dorothea Regulon^63^. The VIPER algorithm calculates the enrichment of gene expression signatures of regulons, that are based on transcription regulatory networks. This way the activity of a transcription factor (TF) is inferred based on the expression of downstream genes known to be regulated by this specific TF. The Dorothea regulon contains known regulatory interactions, annotated based on the confidence that this interaction exists. Here interactions are restricted to confidence levels A and B.

### Hardware and software specifications

All models were expressed in and trained using the PyTorch framework^64^ (version 1.12) in Python (version 3.8.8). When using the 978 landmark genes and for the serology case study, the models were trained in an NVIDIA GeForce RTX 3060 Laptop GPU with 6 GB of memory. The larger models (using 10,086 genes and the single-cell lung fibrosis data) were trained on the MIT Satori GPU cluster using NVIDIA V100 32GB memory GPU cards. Pre-processing and statistical analysis of the results were done in the R programming language (version 4.1.2). Visualization of results was done mainly using *ggplot2*^65^. More information about the versions of each library used can be found in the GitHub provided in the Data and code availability section.

### Data and code availability

The study did not produce new experimental data. All analyzed data that were used to train our models and produce all tables and figures, as well as all the code to generate those data, figures and tables are available at https://github.com/NickMeim/OmicTranslationBenchmark.

## Supporting information

manuscript_neural_net_translation_supplementary

## Acknowledgments

The authors would like to thank Brian Joughin for his valuable input on this work. We acknowledge funding from MIT-IBM Watson AI Lab, National Institutes of Health (NIH) IMPAcTB contract #75N93019C00071, NIH grant U01-AI67892, US Army Research Office Cooperative Agreement W911NF-19-2-0026, and the Swedish Research Council 06349 (AN).

## Author contributions

NM and DAL conceived the study together with input from AN who run a pilot simulation. TNH and SM provided feedback on the methods and implementation of the approach. NM implemented the code and executed the simulations, preprocessed the data, trained the final models and analyzed their results. KP performed downstream analysis for the serology case study and interpreted the results. KP also wrote part of the respective results section. DYZ preprocessed and retrieved the data for the single-cell lung fibrosis case study and helped with result interpretation. NM wrote the manuscript and generated the figures. KP, DYZ, AN, TNH, SM and DAL edited the manuscript.

