## Supplementary material for "Autoencoder Model for Translating Omics Signatures": manuscript_neural_net_translation_supplementary

#### Supplementary Figures

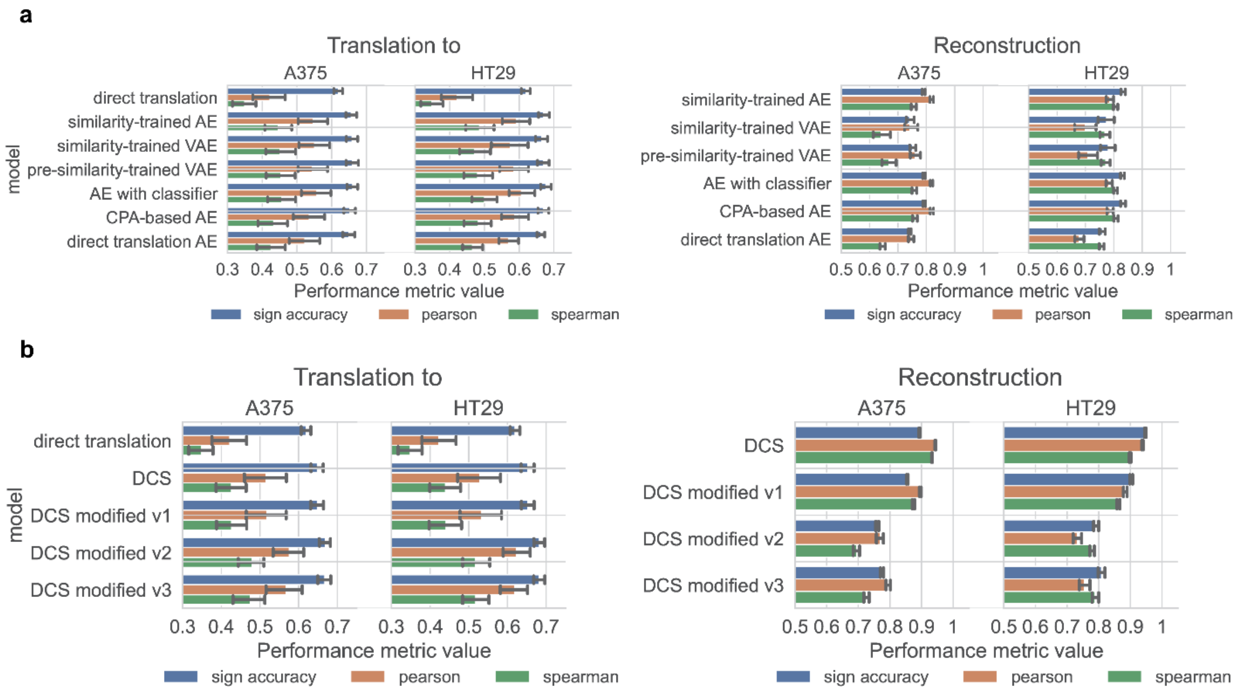


**Supplementary Figure 1:** **a)** Comparisons of variations of our framework when using the 10,086 genes of the L1000 **b)** Comparisons of variations of the DeepCellState model when using the 10,086 genes of the L1000

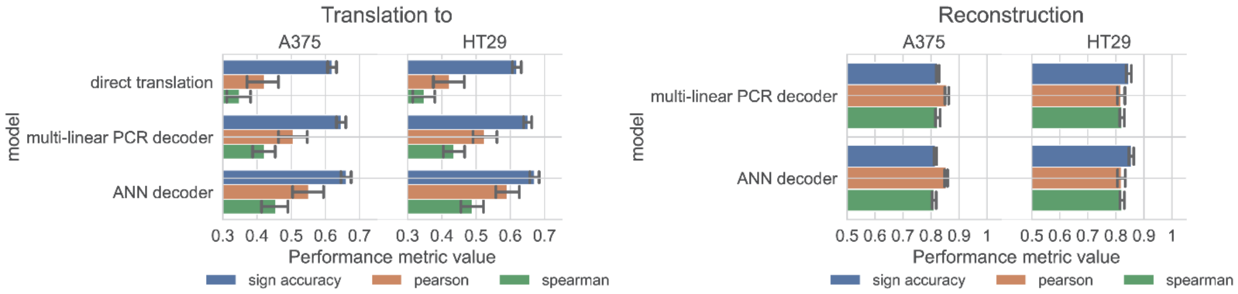


**Supplementary Figure 2:** Comparisons of variations of the TransCompR-based model when using the 10,086 genes of the L1000

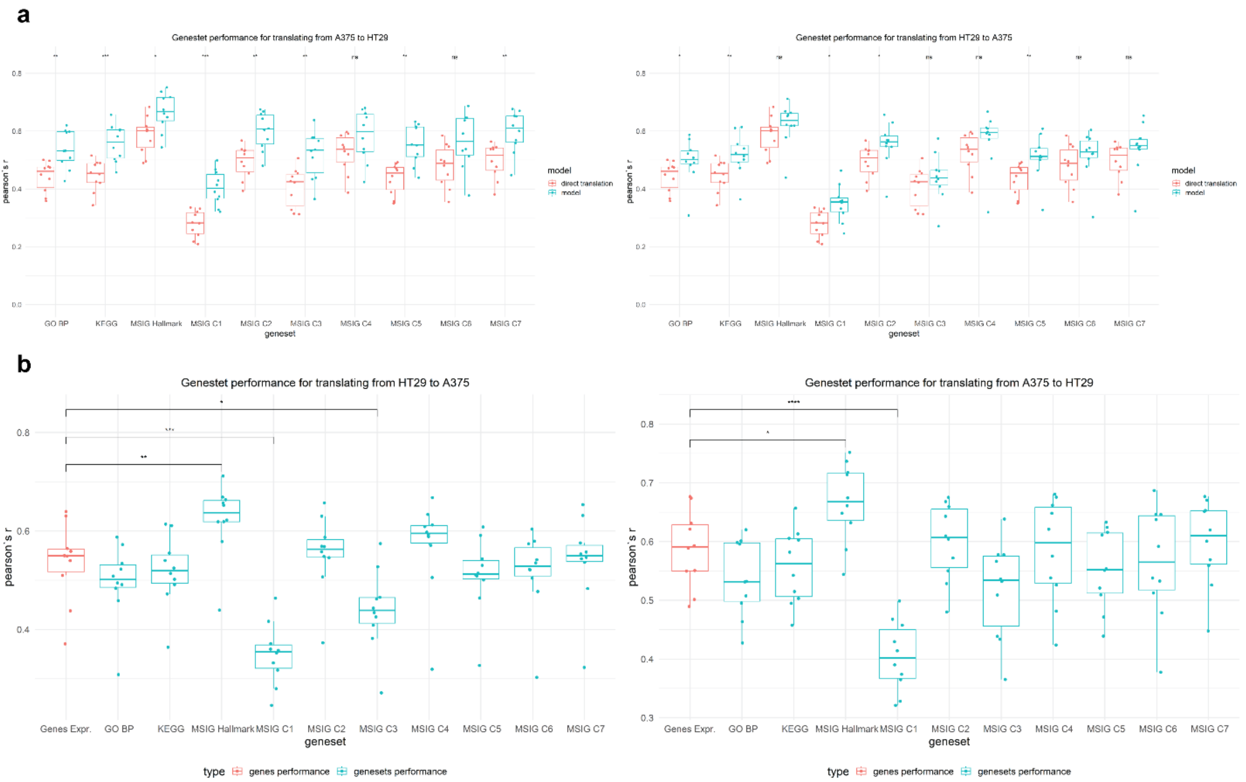


**Supplementary Figure 3:** Performance in inferring enrichment scores for different types of gene sets by using the predicted gene expression. **a)** Comparing the performance of predicted GSEA scores from translated signatures and the performance of direct translation of GSEA scores into another cell line. **b)** Comparison of GSEA scores performance and gene expression translation.

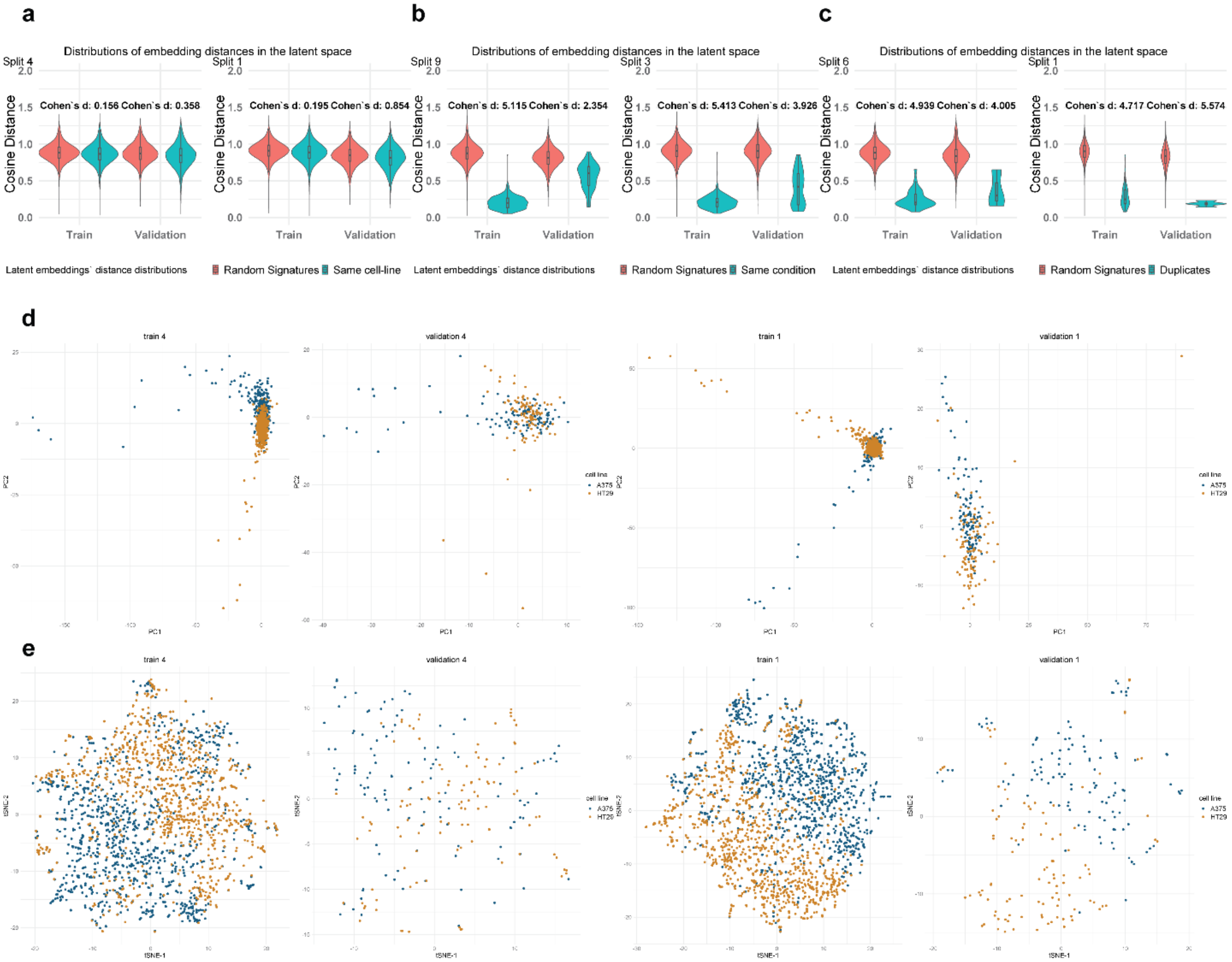


**Supplementary Figure 4:** The embeddings used here are derived from using all the 10,086 L1000 genes with one global latent space, with a jointly trained classifier. **a)** Cosine distance between embeddings coming from random pairs of signatures and pairs coming from the same cell line. **b)** Cosine distance between embeddings coming from random pairs of signatures and pairs coming from the same condition **c)** Cosine distance between embeddings coming from random pairs of signatures and pairs being biological duplicates **d)** PCA visualization for the splits with maximum and minimum separation **e)** t-SNE visualization for the splits with the maximum and minimum separation

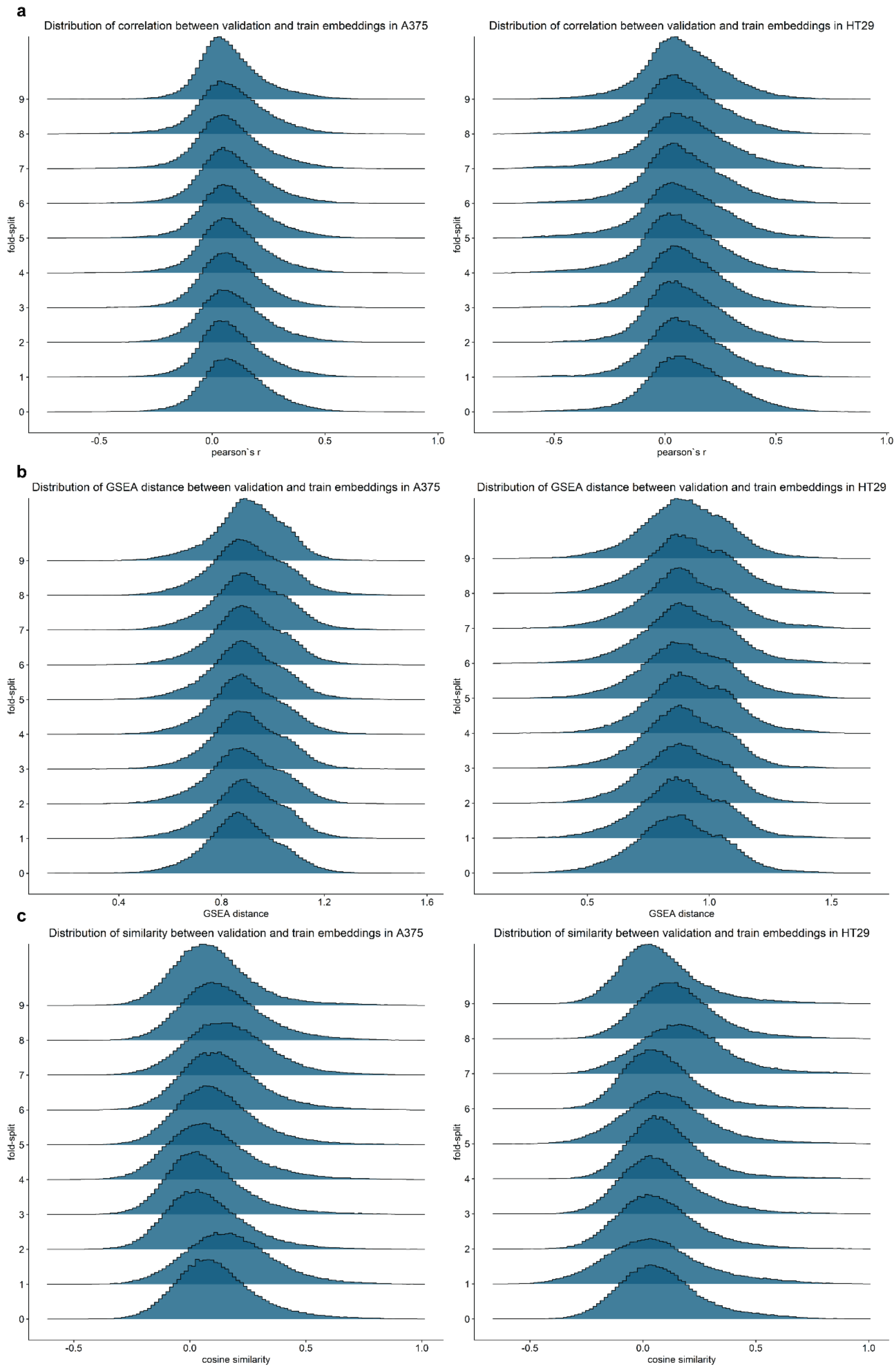


**Supplementary Figure 5:** Similarity of validation samples and training samples in 10-fold cross-validation. **a-b)** Gene expression correlation distribution, between every validation sample and training sample across folds, in A375 and HT29. **c-d)** GSEA-based distance (see Supplementary Method 5) distribution between gene expression profiles of validation and train samples in A375 and HT29. **e-f)** Cosine similarity between latent space embeddings of validation and train samples in A375 and HT29

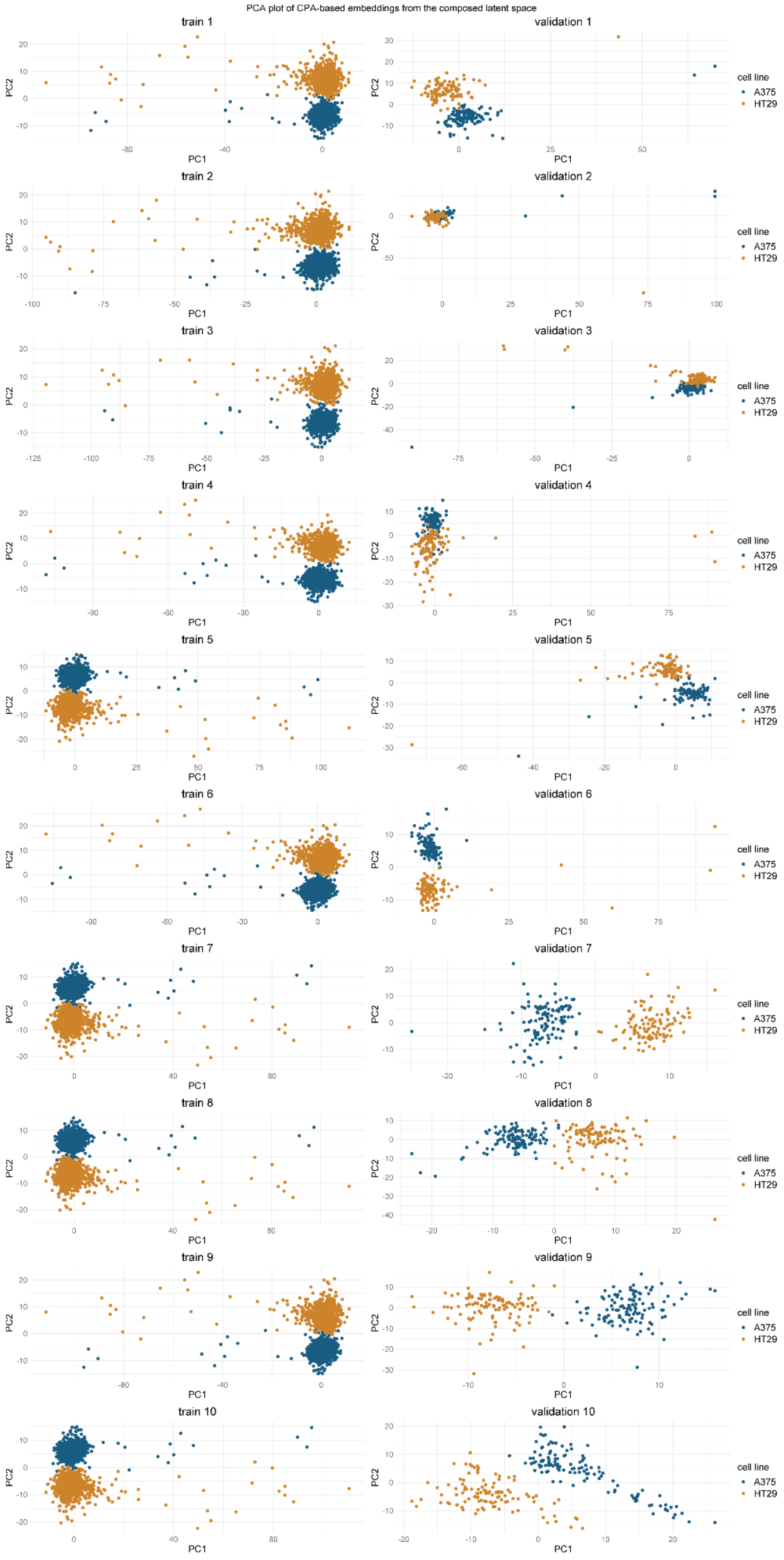


**<u>Supplementary Figure 6:</u>** Composed latent space PCA visualization for embeddings derived from the CPA-based approach using 10,086 L1000 genes.

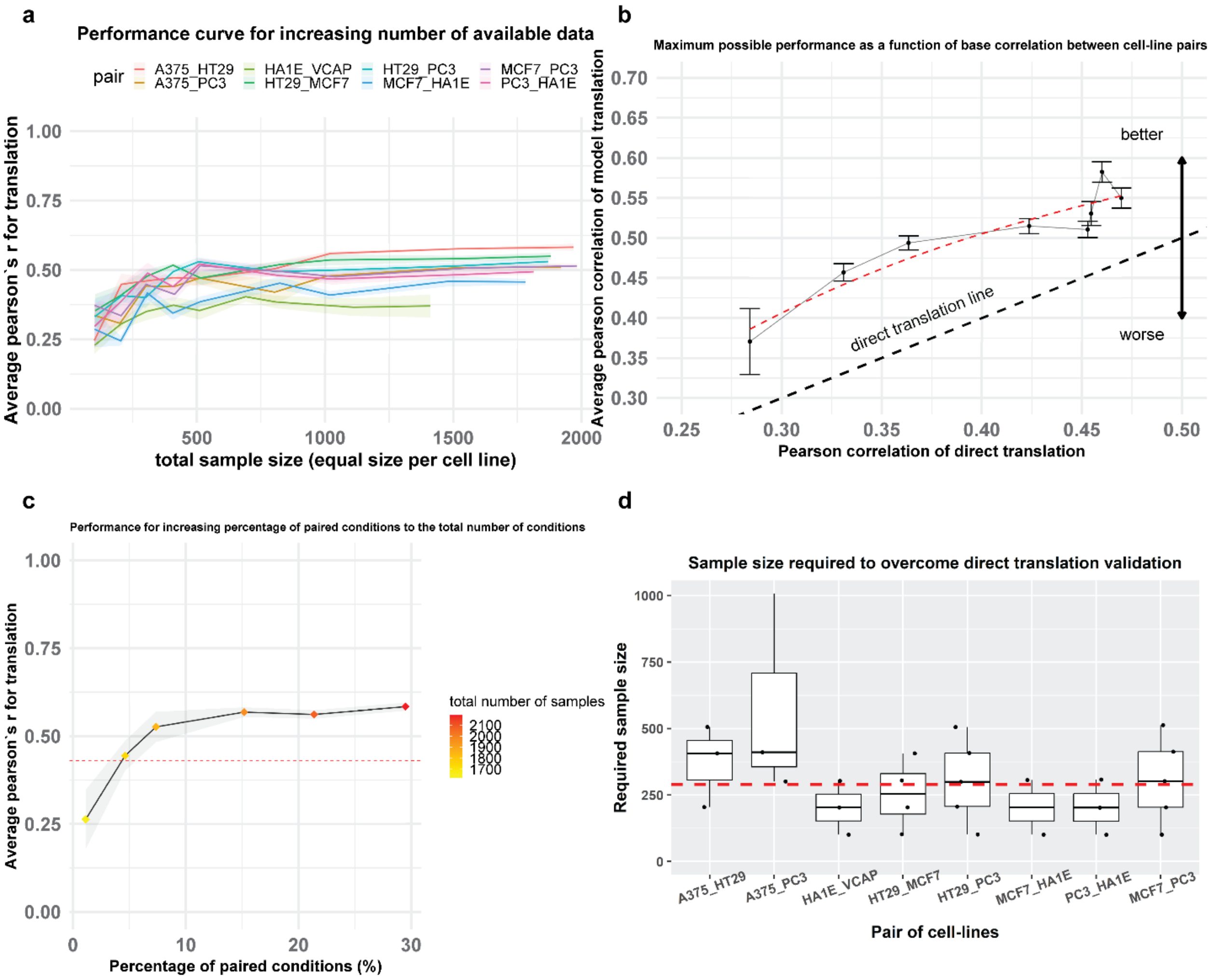


**<u>Supplementary Figure 7:</u>** Performance analysis using all the 10,086 genes for the L1000 **a)** Performance in the translation task of the CPA-based approach across different cell-line pairs and different sizes of training data. **b)** Model performance in translation as a function of the initial similarity of 2 cell lines. **c)** Model performance in translation for different percentages of paired conditions. **d)** Required sample size to overcome direct translation's performance.

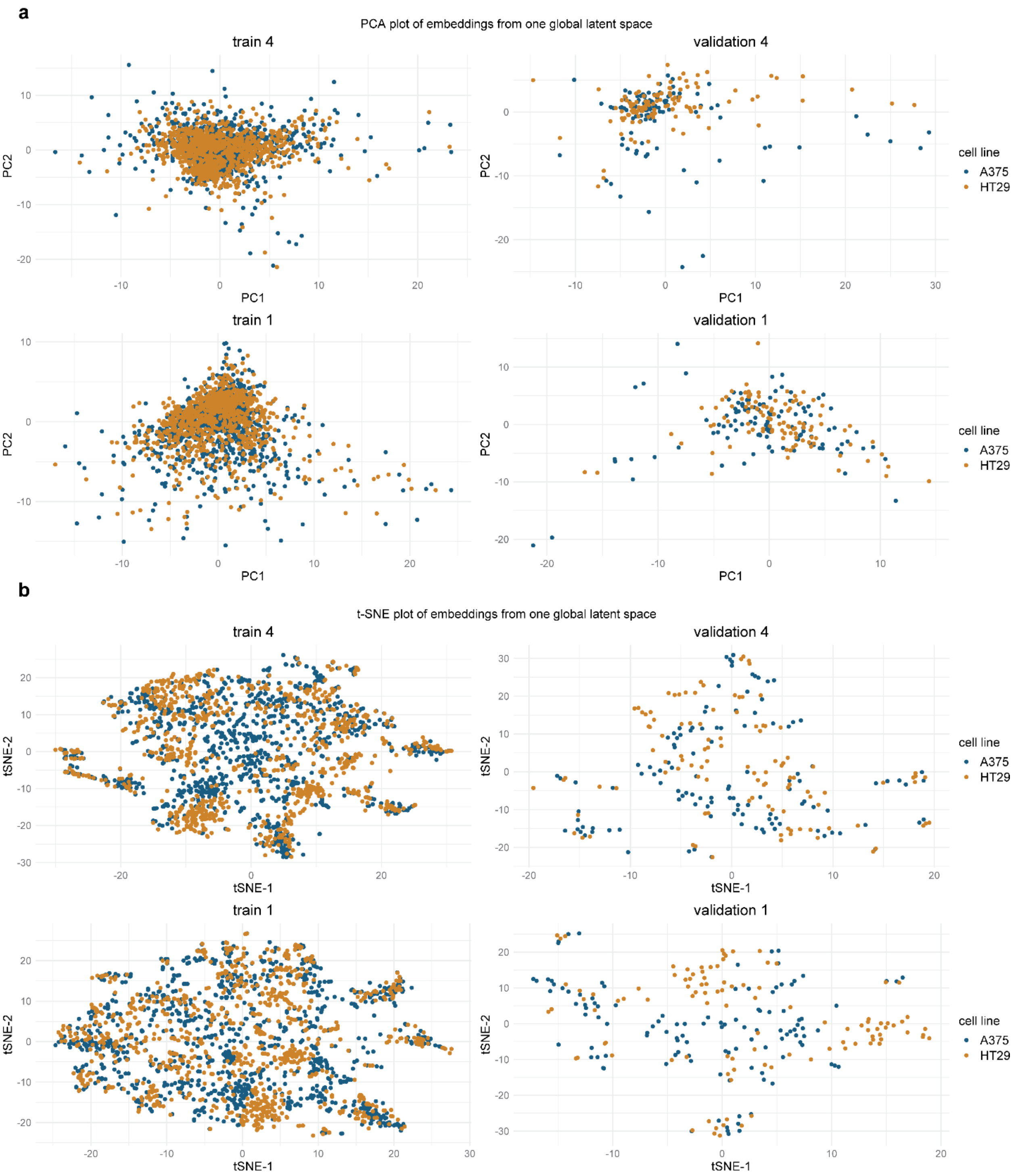


**<u>Supplementary Figure 8:</u>** The two splits in 10-fold cross-validation that are shown each time here are the ones where the maximum and minimum difference between the two distributions is observed. For every other split, the difference is between these two extreme cases. Latent space visualization for embeddings derived from the approach with one global latent space using the 978 landmark genes. **a)** PCA plot of global latent space embeddings **b)** t-SNE plot of global latent space embeddings

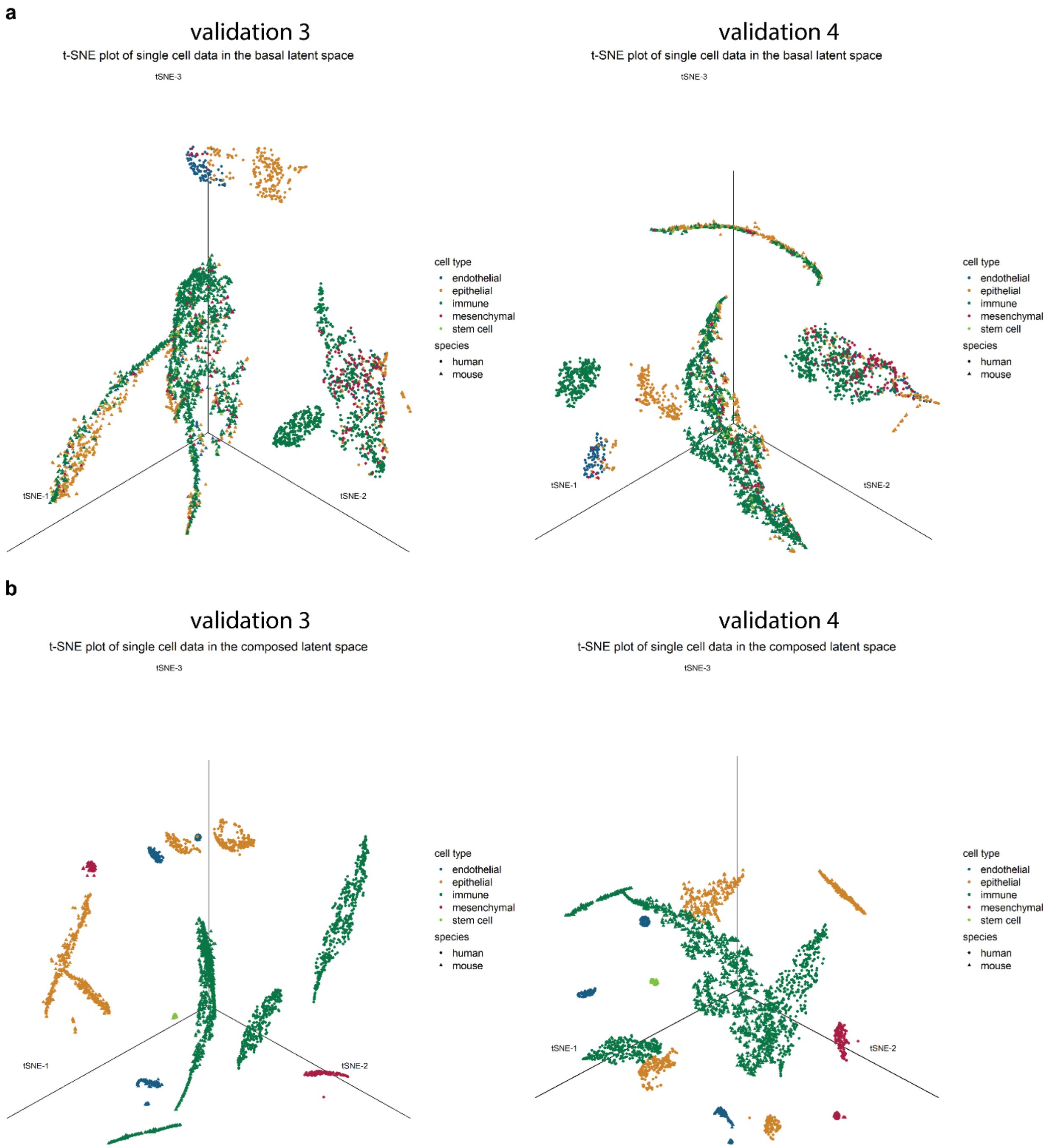


**<u>Supplementary Figure 9:</u>** The two splits in 10-fold cross-validation that are shown each time here are the ones where the maximum and minimum difference between the two distributions is observed. For every other split, the difference is between these two extreme cases. Latent space t-SNE visualization for embeddings derived from lung fibrosis datasets **a)** t-SNE plot of global latent space embeddings **b)** t-SNE plot of composed latent space embeddings

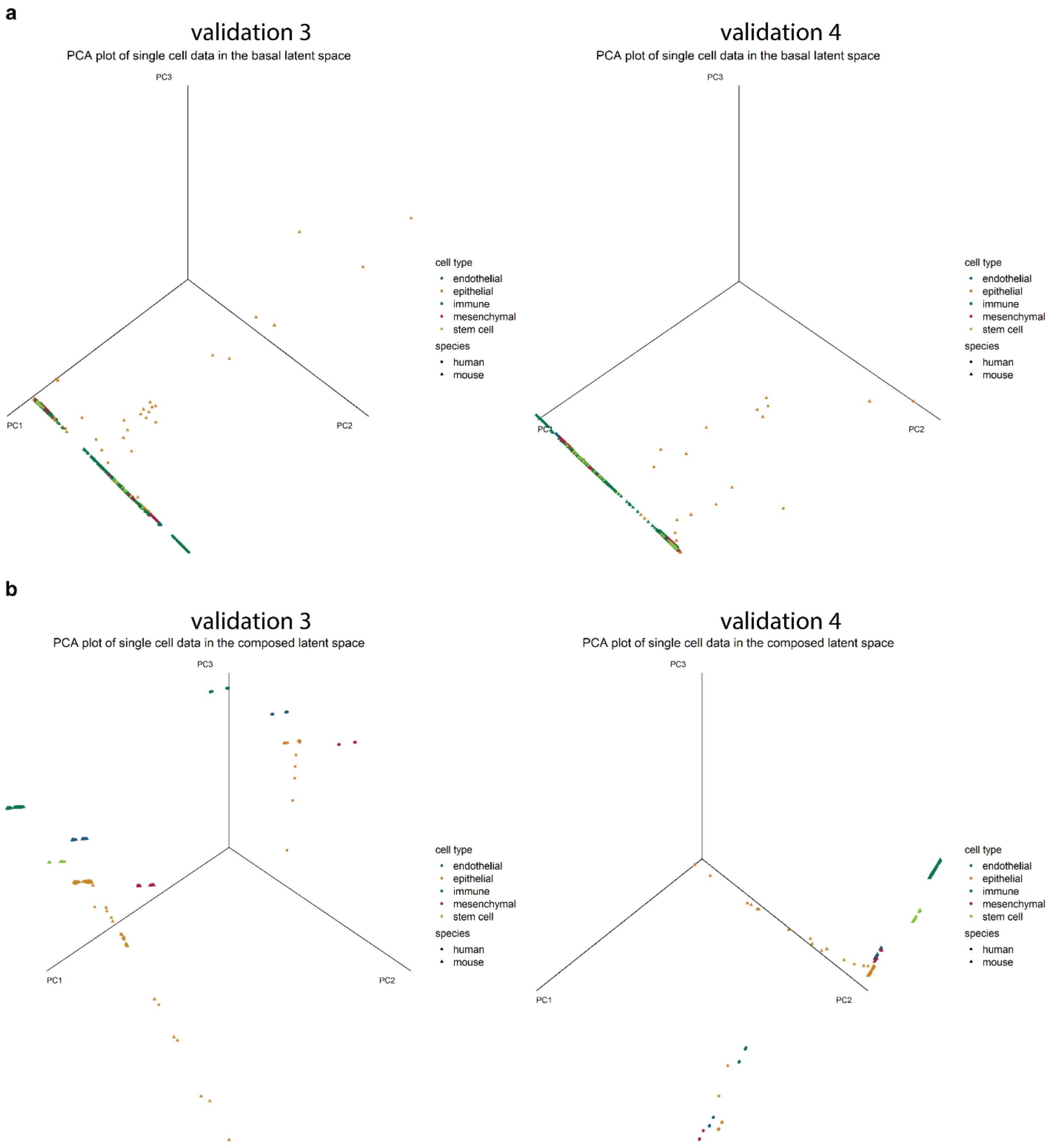


**<u>Supplementary Figure 10:</u>** The two splits in 10-fold cross-validation that are shown each time here are the ones where the maximum and minimum difference between the two distributions is observed. For every other split, the difference is between these two extreme cases. Latent space PCA visualization for embeddings derived from lung fibrosis datasets **a)** PCA plot of global latent space embeddings **b)** PCA plot of composed latent space embeddings

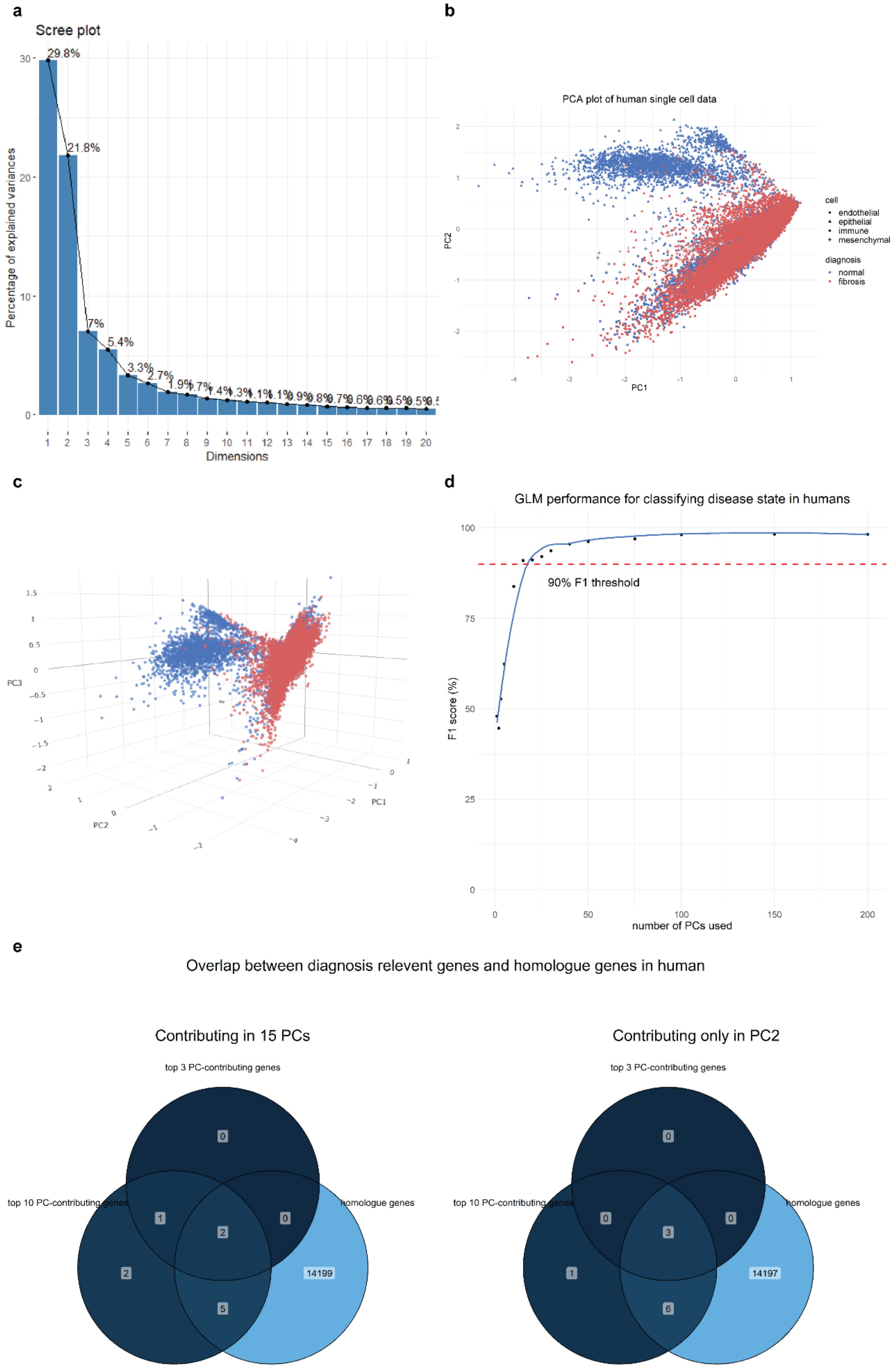


**<u>Supplementary Figure 11:</u>** Important genes analysis for human lung fibrosis **a)** Scree plot showing explained variance of each principal component (PC). The first 3 PCs capture 58.6% of the variance **b-c)**

2-D and 3-D visualization of gene expression data from the human lung fibrosis dataset. The first 3 PCs can separate samples based on fibrosis **d)** Performance in predicting fibrosis of a simple generalized linear model, using different numbers of PCs. Just with 15 PCs 90% F1 score is achieved and saturation of performance begins **e)** Overlap of important genes for human fibrosis, according to their loadings in first PCs, and homolog genes

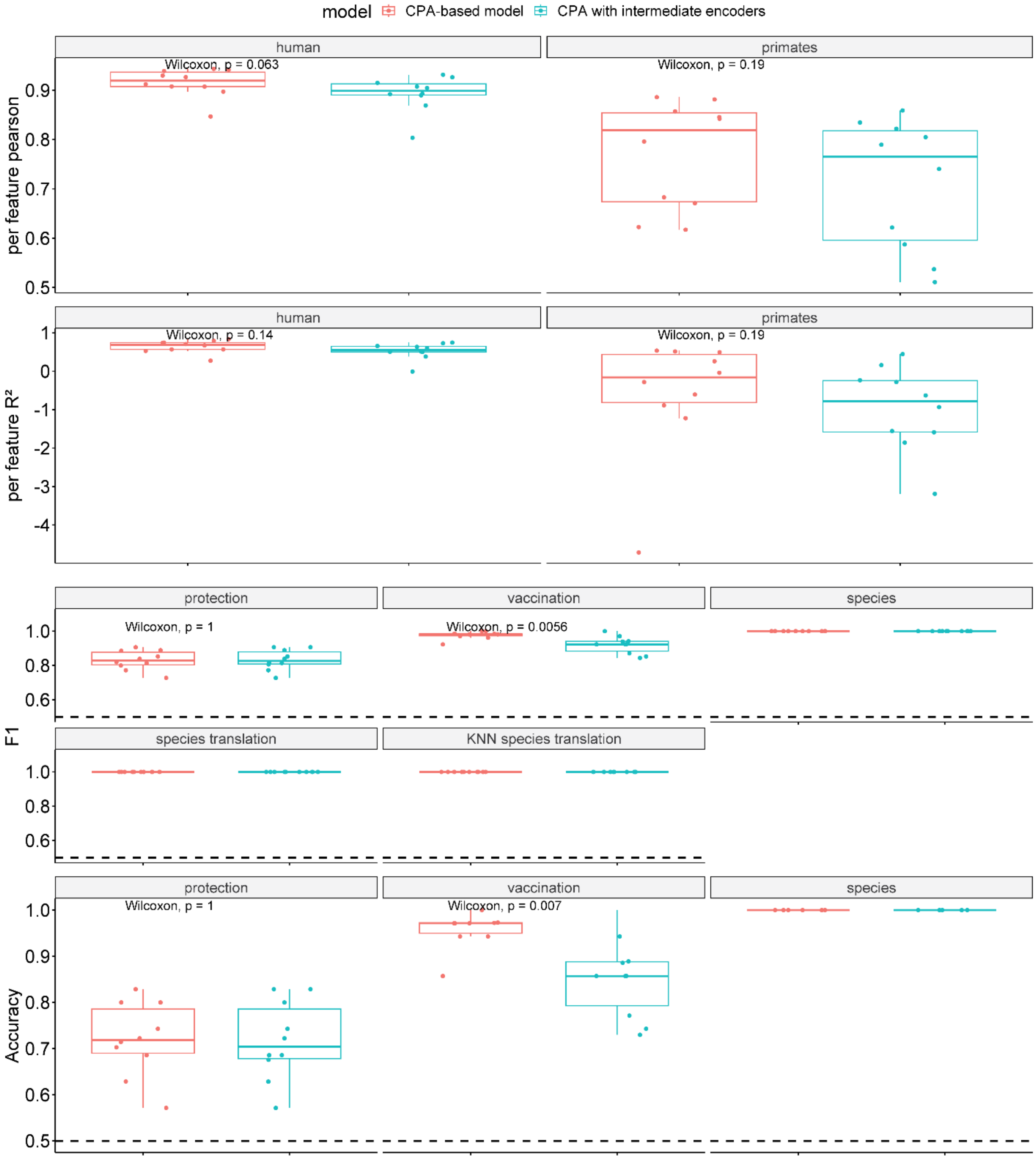


**<u>Supplementary Figure 12</u>** Performance comparison between having a trainable vector or simple ANN to add cell line effect in the serology dataset

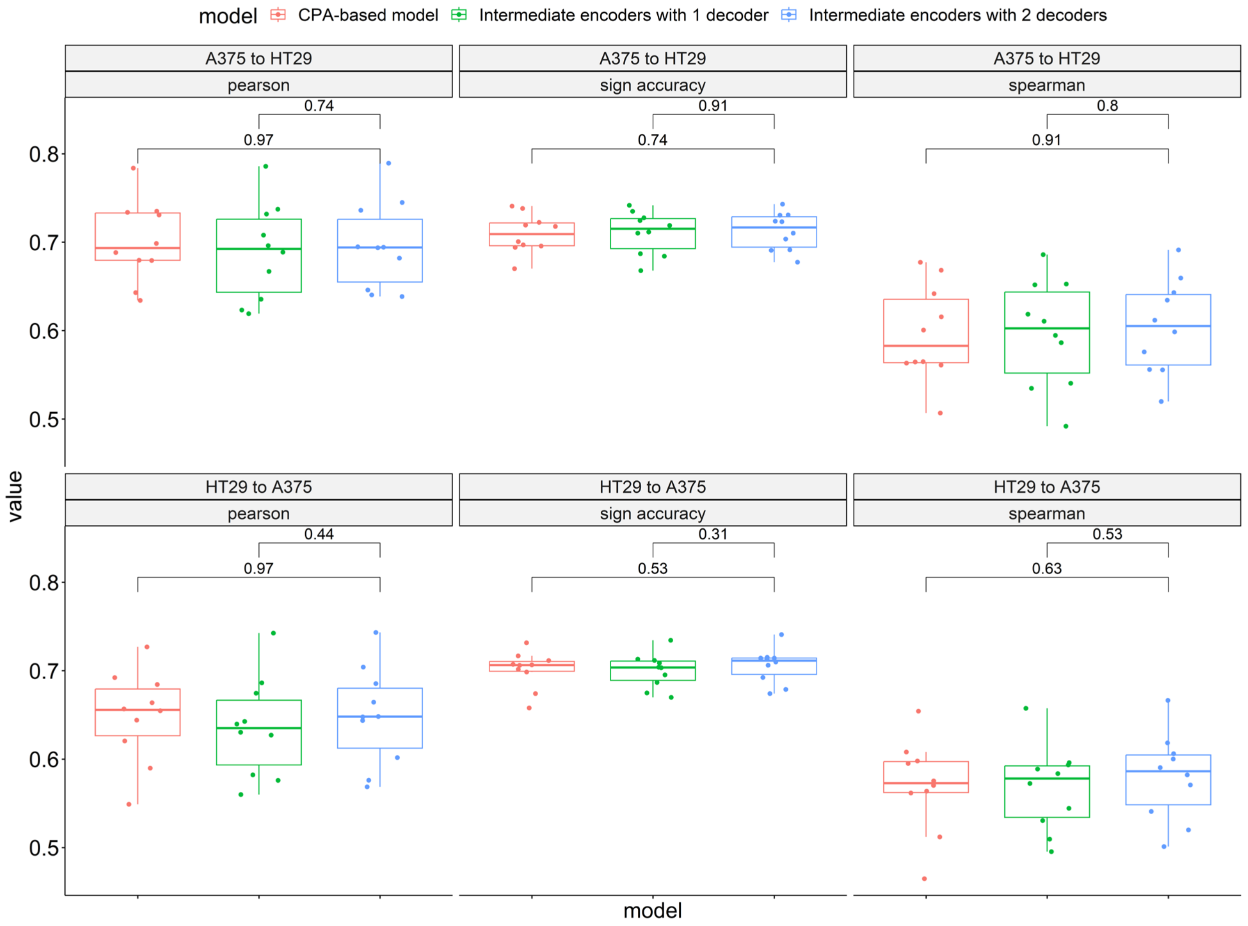


**Supplementary Figure 13:** Performance comparison between having a trainable vector or simple ANN to add cell line effect in the L1000 dataset. The comparison is done in the task of translating the gene expression profile of paired conditions.

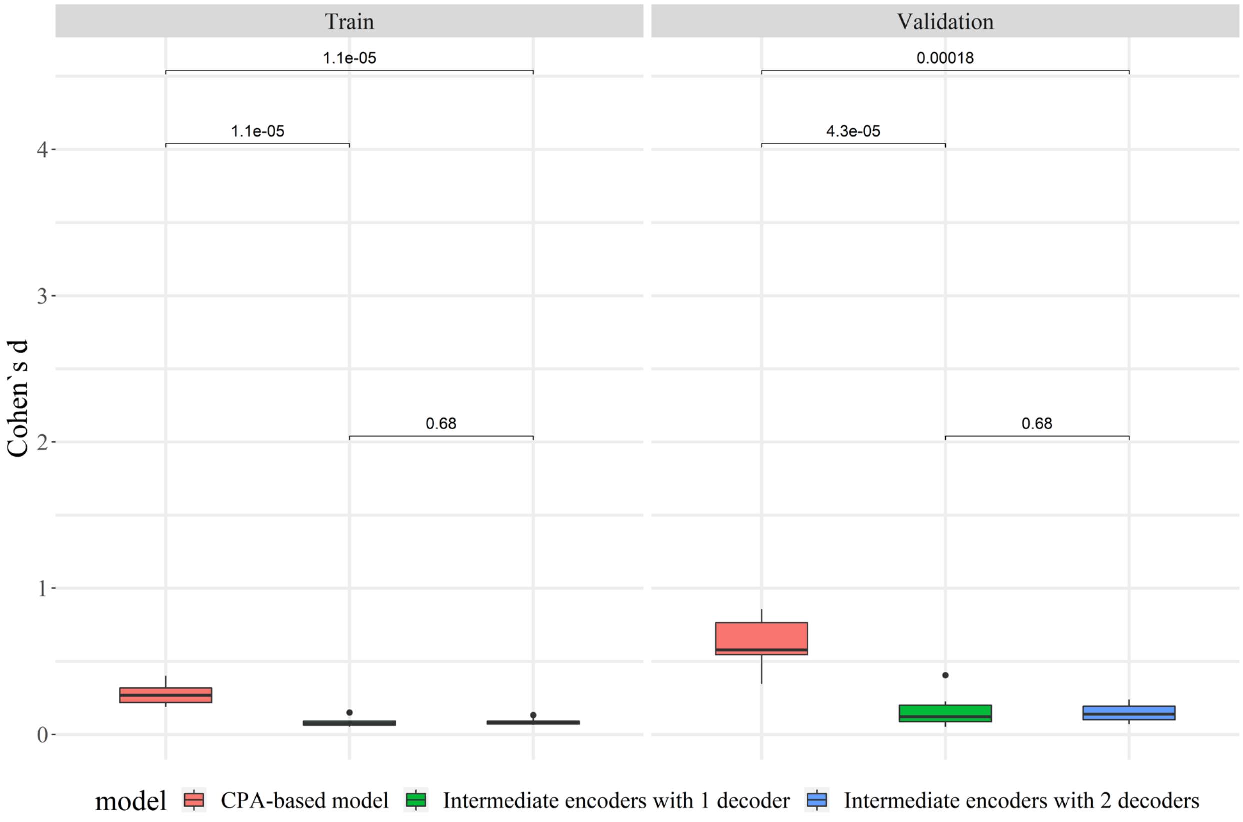


**Supplementary Figure 14:** Comparing Cohen's d between global and composed latent space when using a trainable vector or simple ANN to add cell line effect in the L1000 dataset. Using a trainable vector creates an even bigger difference between composed and basal latent space.

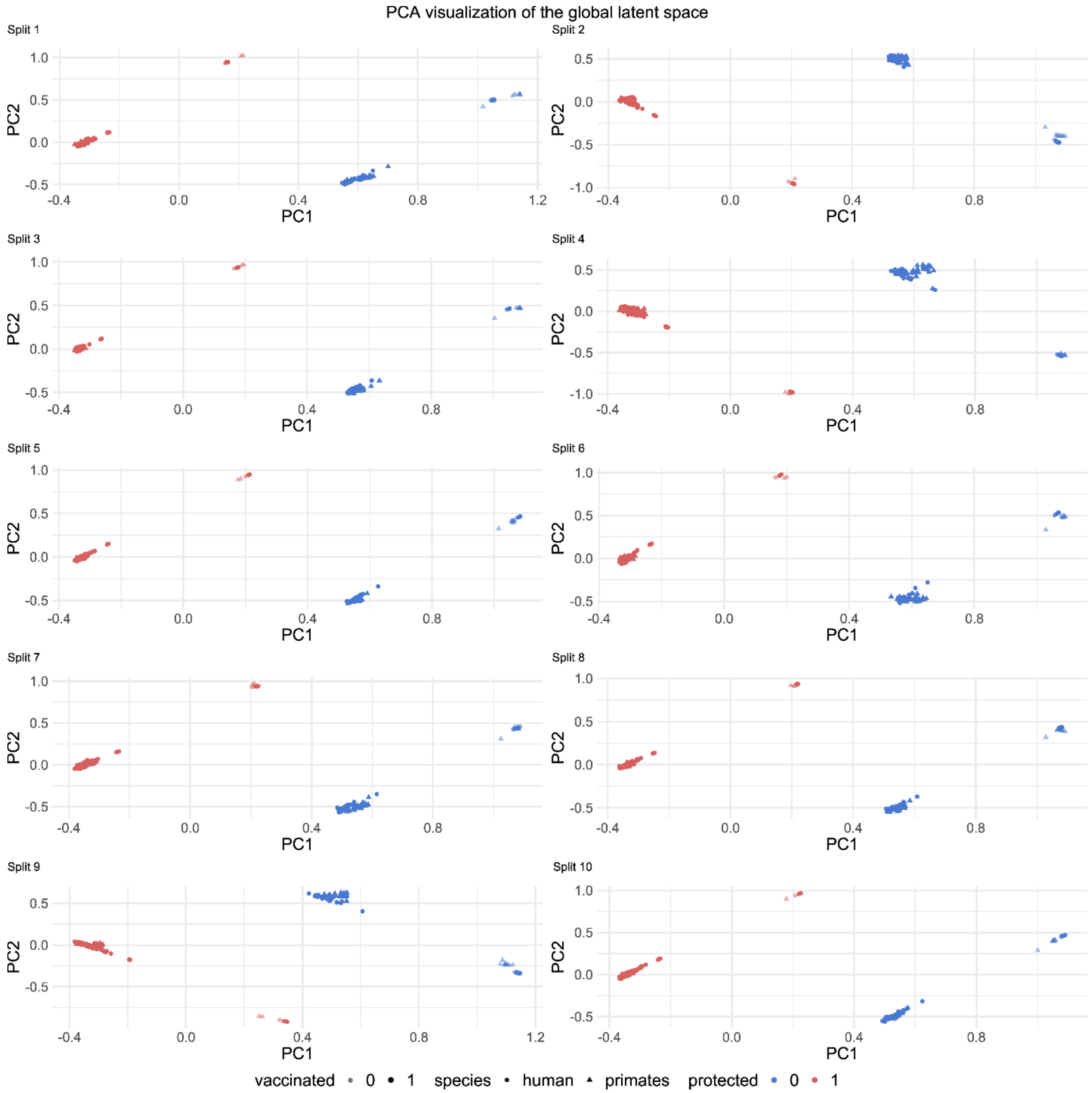


**Supplementary Figure 15:** Global Latent space PCA visualization for embeddings derived from the serology datasets. A clear separation based on protection and vaccination can be observed, while there is no separation based on species

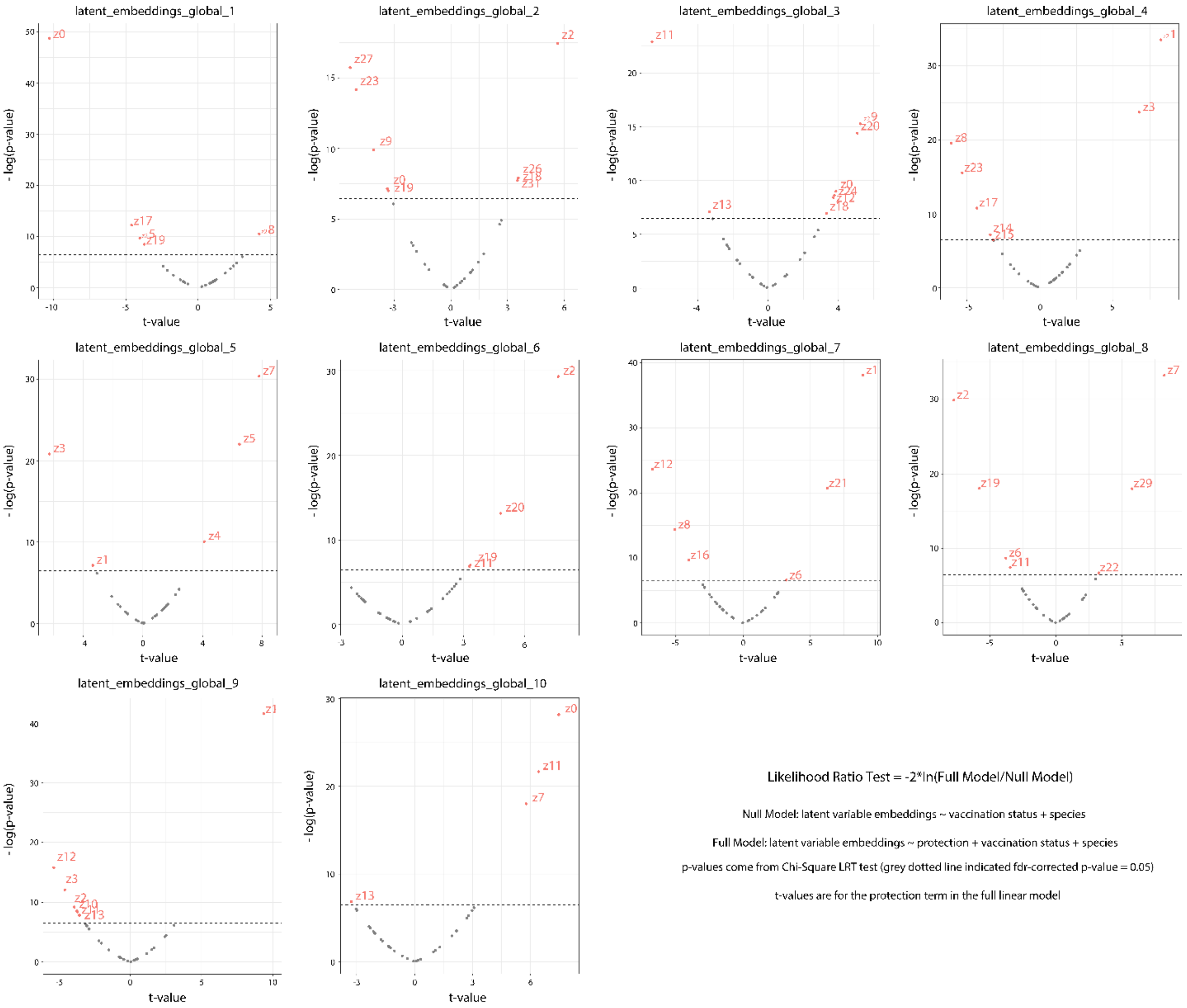


**<u>Supplementary Figure 16:</u>** Visualization of all likelihood ratio tests results across folds, for identifying latent variables associated with viral protection in the serology case study.

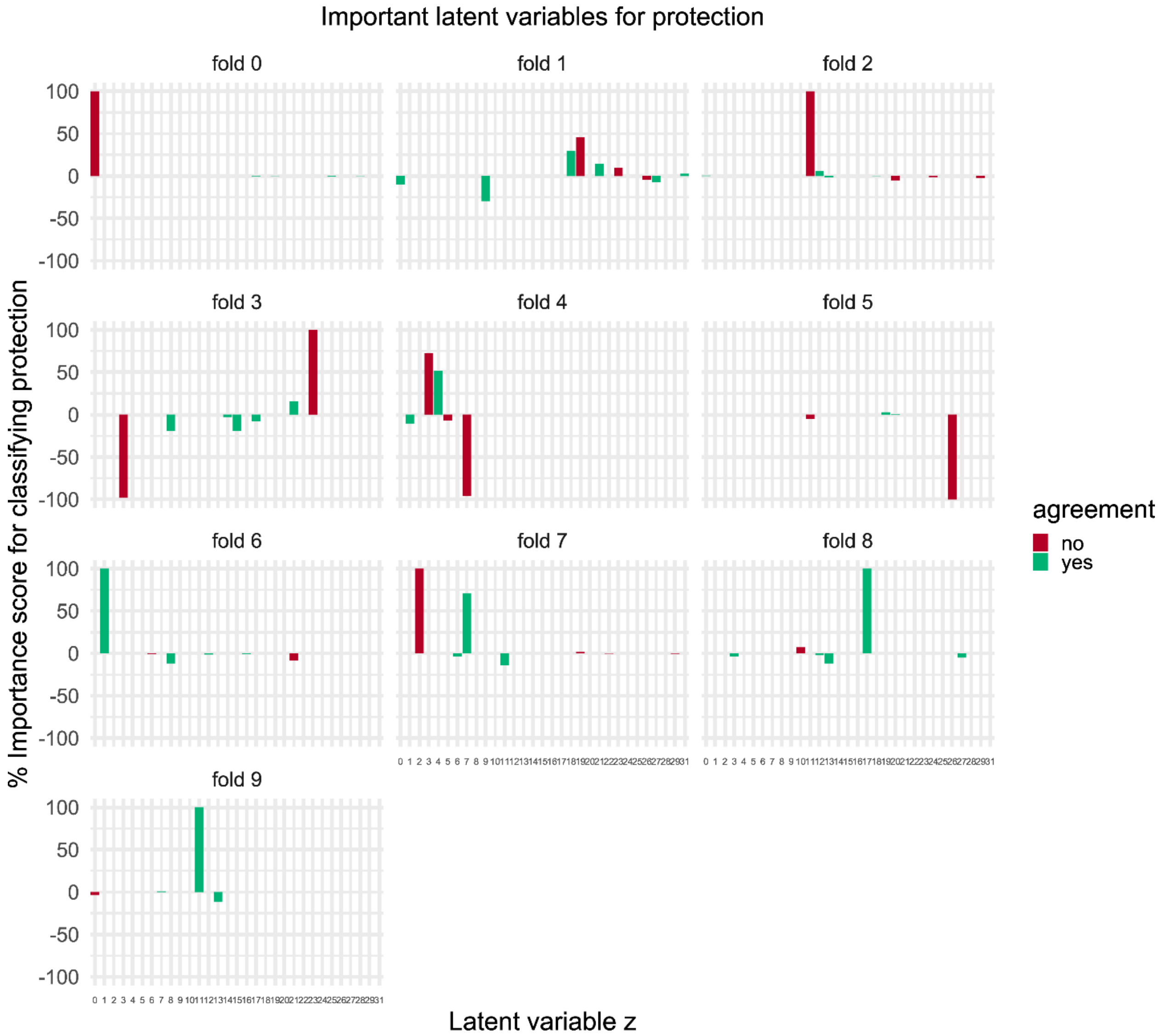


**Supplementary Figure 17:** Importance scores for viral protection of global latent variables, according to the classifier. Only values for statistically significant variables (according to the LRT test are shown). The agreement signifies the agreement in the sign of correlation with protection, between the classifier's score and the score from LRT.

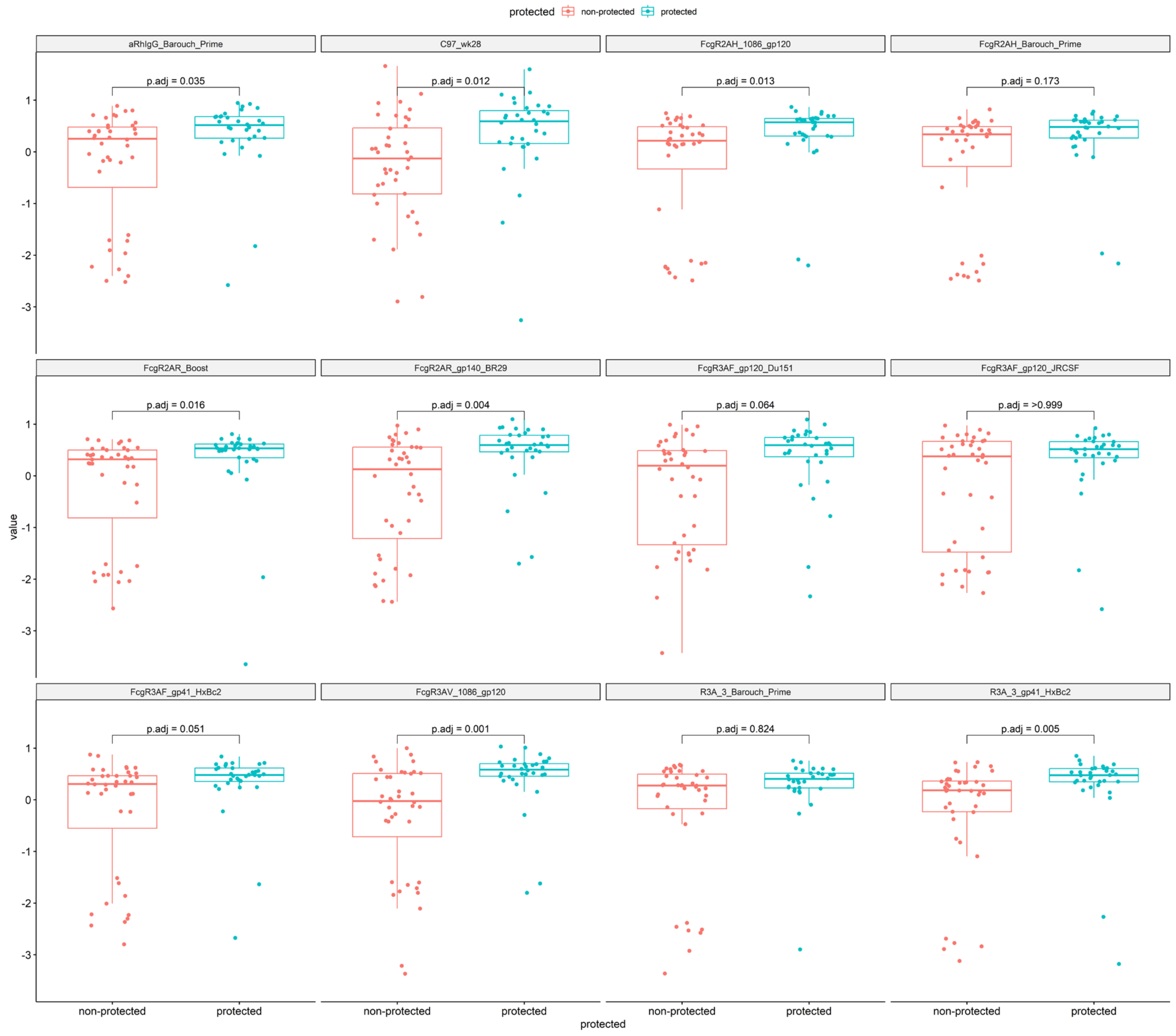


**<u>Supplementary Figure 18:</u>** Distributions of features between protected and non-protected non-human primates. The features shown are those predictive of human viral protection.

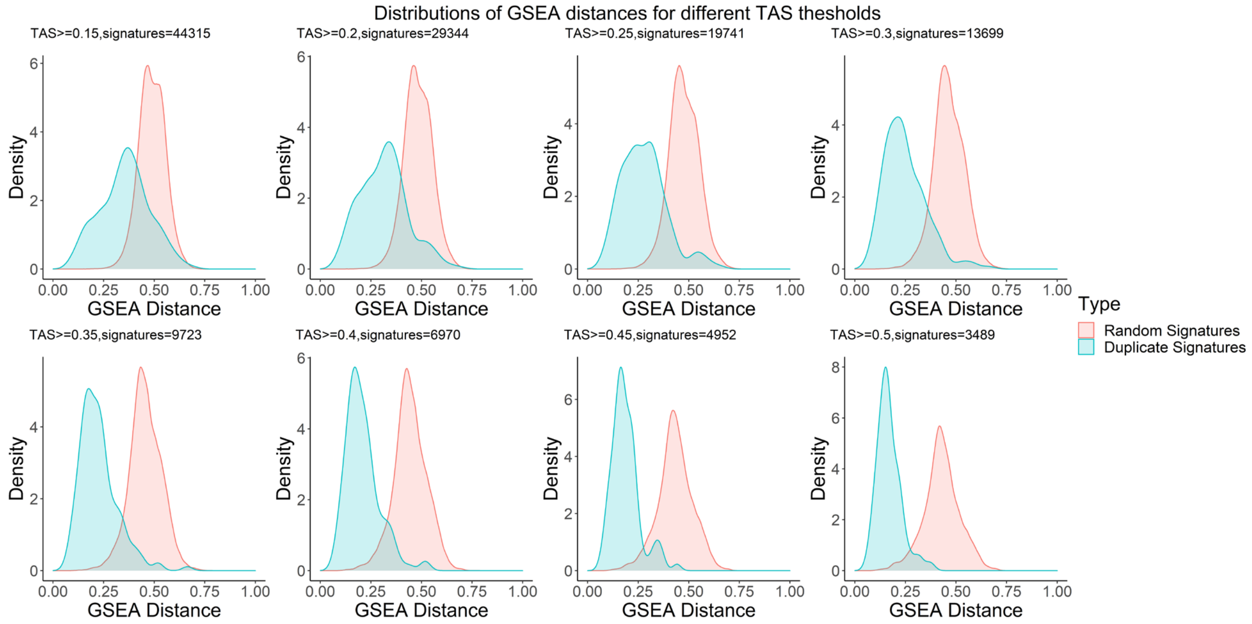


**Supplementary Figure 19:** Separation of GSEA-based distance (Supplementary Methods 5) distributions of gene expression between biological duplicates and randomly selected signatures for varying thresholds of Transcription Activity Score (TAS, see Main Methods), in the case of compound treatment data. There is a statistically significant separation for most TAS thresholds and a very strong separation after TAS>=0.3.

### Supplementary Methods

#### 1. Gene Set Enrichment Analysis (GSEA)

Gene Set Enrichment Analysis (GSEA) was performed on multiple gene sets (Supplementary Figure 3) using the FGSEA library[1,2] from the Bioconductor resource[3]. Thus, the gene-level feature vector of each perturbation was transformed into a gene set-level feature vector of Normalized Enrichment Scores (NES).

#### 2. DeepCellState method and variations

**2.1. Original method:** The original model, as proposed by Umarov, Li, and Arner[4], is an autoencoder neural network framework, which consists of one common encoder and two separate decoders, one for each cell line or species in our case. The model aims to encode every gene expression profile into a common cell line space. The input gene expression is first passed through a dropout layer with a dropout rate of 0.5 and then the encoder consists of fully-connected feedforward neural network layers. The decoders are similar and consist of fully-connected feedforward neural network layers that reconstruct the input gene expression using the latent space representation. The output layer has a direct connection to the dropout layer in the input and combines the two representations to make the final prediction. The authors utilized L1 regularization for the latent layer, enforcing sparsity on the activity of the latent representations. The Activation function used is leaky relu for all layers except the output layer, which uses tanh activation.

**2.2. DCS modified v1:** The model is identical to the originally proposed model, with the only modification of removing the direct connection with the dropout layer in the input.

**2.3. DCS modified v2:** In this variation, we also removed the direct connection with the dropout layer in the input. An important modification is made in the training loss of this model. We include a distance term in the loss, to minimize the distance of latent embeddings coming from the same condition, regardless of the cell or species they are derived from. Finally, for these paired conditions we also minimize the mean squared error of predicted translated gene expression and the ground truth.

**2.4. DCS modified v3:** This variation is identical to version 2, with the only difference of not calculating the mean squared error of direct translation when using the model. We still use a distance term in the loss function.

#### 3. TransCompR-based method

"Translatable Components Regression"[5] (TransCompR) is a method that can map human data into the principal component space of another species to identify translatable animal features that can predict human disease processes and phenotypes. For translating molecular profiles, we use this framework for projecting the molecular profile of a biological system or species into the principal component space of another system or species. This principal component space is now equivalent to the latent space which can be used by a neural network (like the decoder) or a simple multi-linear regression model to predict the translated molecular profile (Supplementary Figure 2).

#### 4. FIT-based method

FIT[6] is a machine learning method that fits a linear regression model between homolog genes coming from the same perturbation tested on two different species (or it can be used with cell lines). During fitting a regularization penalty is added to force the slope of the fitted line to be 1 and the intercept 0. This trained framework can be used then to translate molecular profiles.

#### 5. GSEA-based distance of transcriptomic profiles

The pairwise distance between gene expression feature vectors was calculated using the R package Gene Expression Signature in Bioconductor[7], similar to Iorio et al.[8] Given two gene expression vectors ranked by their z-scored expression, A and B, GSEA is used to calculate the ES of the top and bottom genes of A in B and vice versa. The distance between the gene expression profiles is computed as $1 - \frac{ES_{A\ in\ B} + ES_{B\ in\ A}}{2}$ and ranges from 0 to 2. A GSEA distance equal to 0 means that the most upregulated and downregulated genes are the same in the two vectors A and B, while a distance equal to 2 means they are reversed. The GSEA distance is calculated for multiple thresholds as to how many top and bottom genes to consider and the average distance is taken for further analysis.

### Supplementary Tables

**Table 1:** P-values from comparing methods from Figure 1b in the task of translation. The Mann–Whitney U test was used to compare the performance of the different methods

| | | A375 | | | | | | | | | HT29 | | | | | | | | |
|---|---|---|---|---|---|---|---|---|---|---|---|---|---|---|---|---|---|---|---|
| | | Autoencoders with classifier | CPA-based Autoencoders | DCS | DCS modified v1 | DCS modified v2 | FIT | TransCompR | direct translation | similarity-trained Autoencoders | Autoencoders with classifier | CPA-based Autoencoders | DCS | DCS modified v1 | DCS modified v2 | FIT | TransCompR | direct translation | similarity-trained Autoencoders |
| Autoencoders with classifier | pearson | 1 | 0.6232 | | | | 0.0013 | 0.3847 | | | 1 | 0.7337 | | | | $\leq 10^{-3}$ | 0.3075 | | |
| | sign accuracy | 1 | 0.4727 | | | | $\leq 10^{-3}$ | 0.1212 | | | 1 | 0.6776 | | | | $\leq 10^{-3}$ | 0.1405 | | |
| | spearman | 1 | 0.5205 | | | | $\leq 10^{-3}$ | 0.1859 | | | 1 | 0.6776 | | | | 0.014 | 0.2123 | | |
| CPA-based Autoencoders | pearson | | 1 | | | | 0.0017 | 0.6232 | | | | 1 | | | | | | | |
| | sign accuracy | | 1 | | | | $\leq 10^{-3}$ | 0.1212 | | | | 1 | | | | | | | |
| | spearman | | 1 | | | | $\leq 10^{-3}$ | 0.3075 | | | | 1 | | | | | | | |
| DCS | pearson | $\leq 10^{-3}$ | $\leq 10^{-3}$ | 1 | 0.0036 | $\leq 10^{-3}$ | 0.0539 | $\leq 10^{-3}$ | 0.089 | $\leq 10^{-3}$ | $\leq 10^{-3}$ | $\leq 10^{-3}$ | 1 | 0.0022 | $\leq 10^{-3}$ | 0.0312 | $\leq 10^{-3}$ | 0.0757 | $\leq 10^{-3}$ |
| | sign accuracy | $\leq 10^{-3}$ | 0.0013 | 1 | 0.0058 | $\leq 10^{-3}$ | $\leq 10^{-3}$ | 0.0013 | 0.064 | $\leq 10^{-3}$ | $\leq 10^{-3}$ | $\leq 10^{-3}$ | 1 | 0.0036 | $\leq 10^{-3}$ | $\leq 10^{-3}$ | 0.001 | 0.0452 | $\leq 10^{-3}$ |
| | spearman | $\leq 10^{-3}$ | 0.0013 | 1 | 0.0036 | $\leq 10^{-3}$ | $\leq 10^{-3}$ | 0.0013 | 0.0376 | $\leq 10^{-3}$ | $\leq 10^{-3}$ | $\leq 10^{-3}$ | 1 | 0.001 | $\leq 10^{-3}$ | $\leq 10^{-3}$ | $\leq 10^{-3}$ | 0.0312 | $\leq 10^{-3}$ |
| DCS modified v1 | pearson | 0.1212 | 0.2413 | | 1 | 0.2123 | 0.089 | 0.3847 | $\leq 10^{-3}$ | 0.1212 | 0.0312 | 0.0376 | | 1 | 0.0452 | 0.0376 | 0.1859 | $\leq 10^{-3}$ | 0.0257 |
| | sign accuracy | 0.1041 | 0.2413 | | 1 | 0.089 | $\leq 10^{-3}$ | 0.9698 | $\leq 10^{-3}$ | 0.0757 | 0.3075 | 0.273 | | 1 | 0.1405 | $\leq 10^{-3}$ | 0.9698 | $\leq 10^{-3}$ | 0.1405 |
| | spearman | 0.2123 | 0.5205 | | 1 | 0.1859 | $\leq 10^{-3}$ | 1 | $\leq 10^{-3}$ | 0.1859 | 0.2413 | 0.3847 | | 1 | 0.1859 | $\leq 10^{-3}$ | 0.9097 | $\leq 10^{-3}$ | 0.2123 |
| DCS modified v2 | pearson | 0.7337 | 0.9097 | | | 1 | 0.0013 | 0.5708 | $\leq 10^{-3}$ | 0.7337 | 0.7337 | 0.9097 | | | 1 | $\leq 10^{-3}$ | 0.3847 | $\leq 10^{-3}$ | 0.6232 |
| | sign accuracy | 0.9698 | 0.5708 | | | 1 | $\leq 10^{-3}$ | 0.0757 | $\leq 10^{-3}$ | 0.9097 | 0.8501 | 0.5205 | | | 1 | $\leq 10^{-3}$ | 0.0757 | $\leq 10^{-3}$ | 1 |
| | spearman | 0.9097 | 0.3847 | | | 1 | 0.001 | 0.1041 | $\leq 10^{-3}$ | 0.9097 | 0.7913 | 0.4727 | | | 1 | 0.0173 | 0.162 | $\leq 10^{-3}$ | 0.8501 |
| direct translation | pearson | $\leq 10^{-3}$ | $\leq 10^{-3}$ | | | | 0.001 | $\leq 10^{-3}$ | 1 | | $\leq 10^{-3}$ | $\leq 10^{-3}$ | | | | $\leq 10^{-3}$ | $\leq 10^{-3}$ | 1 | |
| | sign accuracy | $\leq 10^{-3}$ | $\leq 10^{-3}$ | | | | $\leq 10^{-3}$ | $\leq 10^{-3}$ | 1 | | $\leq 10^{-3}$ | $\leq 10^{-3}$ | | | | $\leq 10^{-3}$ | $\leq 10^{-3}$ | 1 | |
| | spearman | $\leq 10^{-3}$ | $\leq 10^{-3}$ | | | | $\leq 10^{-3}$ | $\leq 10^{-3}$ | 1 | | $\leq 10^{-3}$ | $\leq 10^{-3}$ | | | | $\leq 10^{-3}$ | $\leq 10^{-3}$ | 1 | |
| FIT | pearson | | | | | | 1 | 0.0046 | | | | $\leq 10^{-3}$ | | | | 1 | $\leq 10^{-3}$ | | |
| | sign accuracy | | | | | | 1 | $\leq 10^{-3}$ | | | | $\leq 10^{-3}$ | | | | 1 | $\leq 10^{-3}$ | | |
| | spearman | | | | | | 1 | $\leq 10^{-3}$ | | | | 0.0058 | | | | 1 | $\leq 10^{-3}$ | | |
| similarity-trained Autoencoders | pearson | 0.8501 | 0.6776 | | | | $\leq 10^{-3}$ | 0.273 | $\leq 10^{-3}$ | 1 | 0.5205 | 0.4727 | | | | $\leq 10^{-3}$ | 0.2123 | $\leq 10^{-3}$ | 1 |
| | sign accuracy | 1 | 0.4727 | | | | $\leq 10^{-3}$ | 0.0452 | $\leq 10^{-3}$ | 1 | 0.9097 | 0.5708 | | | | $\leq 10^{-3}$ | 0.1212 | $\leq 10^{-3}$ | 1 |
| | spearman | 0.9698 | 0.4727 | | | | $\leq 10^{-3}$ | 0.1212 | $\leq 10^{-3}$ | 1 | 0.9097 | 0.4274 | | | | 0.0257 | 0.162 | $\leq 10^{-3}$ | 1 |
| TransCompR | pearson | | | | | | | 1 | | | | 0.4274 | | | | | 1 | | |
| | sign accuracy | | | | | | | 1 | | | | 0.2413 | | | | | 1 | | |
| | spearman | | | | | | | 1 | | | | 0.4727 | | | | | 1 | | |

**Table 2:** P-values from comparing methods from Figure 1b in the task of reconstruction. The Mann–Whitney U test was used to compare the performance of the different methods

| | | A375 | | | | | | | HT29 | | | | | | |
|---|---|---|---|---|---|---|---|---|---|---|---|---|---|---|---|
| | | Autoencoders with classifier | CPA-based Autoencoders | DCS | DCS modified v1 | DCS modified v2 | TransCompR | similarity-trained Autoencoders | Autoencoders with classifier | CPA-based Autoencoders | DCS | DCS modified v1 | DCS modified v2 | TransCompR | similarity-trained Autoencoders |
| Autoencoders with classifier | pearson | 1 | 0.6232 | | | | 0.3847 | | 1 | 0.273 | | | | 1 | |
| | sign accuracy | 1 | 0.2411 | | | | 0.0022 | | 1 | 0.4274 | | | | 0.0376 | |
| | spearman | 1 | 0.2123 | | | | 0.1859 | | 1 | 0.3075 | | | | 0.1212 | |
| CPA-based Autoencoders | pearson | | 1 | | | | | | | 1 | | | | | |
| | sign accuracy | | 1 | | | | | | | 1 | | | | | |
| | spearman | | 1 | | | | | | | 1 | | | | | |
| DCS | pearson | $\leq 10^{-3}$ | $\leq 10^{-3}$ | 1 | 0.0036 | $\leq 10^{-3}$ | $\leq 10^{-3}$ | $\leq 10^{-3}$ | $\leq 10^{-3}$ | $\leq 10^{-3}$ | 1 | $\leq 10^{-3}$ | $\leq 10^{-3}$ | $\leq 10^{-3}$ | $\leq 10^{-3}$ |
| | sign accuracy | $\leq 10^{-3}$ | $\leq 10^{-3}$ | 1 | $\leq 10^{-3}$ | $\leq 10^{-3}$ | $\leq 10^{-3}$ | $\leq 10^{-3}$ | $\leq 10^{-3}$ | $\leq 10^{-3}$ | 1 | $\leq 10^{-3}$ | $\leq 10^{-3}$ | $\leq 10^{-3}$ | $\leq 10^{-3}$ |
| | spearman | $\leq 10^{-3}$ | $\leq 10^{-3}$ | 1 | $\leq 10^{-3}$ | $\leq 10^{-3}$ | $\leq 10^{-3}$ | $\leq 10^{-3}$ | $\leq 10^{-3}$ | $\leq 10^{-3}$ | 1 | $\leq 10^{-3}$ | $\leq 10^{-3}$ | $\leq 10^{-3}$ | $\leq 10^{-3}$ |
| DCS modified v1 | pearson | 0.1212 | 0.2413 | | 1 | 0.2123 | 0.3847 | 0.1212 | 0.0539 | 0.0036 | | 1 | $\leq 10^{-3}$ | 0.064 | 0.0757 |
| | sign accuracy | 0.0113 | 0.0028 | | 1 | $\leq 10^{-3}$ | $\leq 10^{-3}$ | 0.0173 | 0.273 | 0.0757 | | 1 | $\leq 10^{-3}$ | 0.0058 | 0.273 |
| | spearman | 0.0017 | $\leq 10^{-3}$ | | 1 | $\leq 10^{-3}$ | $\leq 10^{-3}$ | 0.0173 | 0.1041 | 0.0312 | | 1 | $\leq 10^{-3}$ | 0.0113 | 0.064 |
| DCS modified v2 | pearson | 0.7337 | 0.9097 | | | 1 | 0.5708 | 0.7337 | $\leq 10^{-3}$ | $\leq 10^{-3}$ | | | 1 | $\leq 10^{-3}$ | $\leq 10^{-3}$ |
| | sign accuracy | $\leq 10^{-3}$ | $\leq 10^{-3}$ | | | 1 | $\leq 10^{-3}$ | $\leq 10^{-3}$ | $\leq 10^{-3}$ | $\leq 10^{-3}$ | | | 1 | $\leq 10^{-3}$ | $\leq 10^{-3}$ |
| | spearman | $\leq 10^{-3}$ | $\leq 10^{-3}$ | | | 1 | $\leq 10^{-3}$ | $\leq 10^{-3}$ | $\leq 10^{-3}$ | $\leq 10^{-3}$ | | | 1 | $\leq 10^{-3}$ | $\leq 10^{-3}$ |
| similarity-trained Autoencoders | pearson | 0.8501 | 0.6776 | | | | 0.273 | 1 | 0.7337 | 0.1859 | | | | 0.6232 | 1 |
| | sign accuracy | 0.7337 | 0.4272 | | | | 0.0036 | 1 | 0.7913 | 0.4274 | | | | 0.0452 | 1 |
| | spearman | 0.9698 | 0.2413 | | | | 0.162 | 1 | 0.9698 | 0.273 | | | | 0.1405 | 1 |
| TransCompR | pearson | | 0.6232 | | | | 1 | | | 0.273 | | | | 1 | |
| | sign accuracy | | 0.0211 | | | | 1 | | | 0.1405 | | | | 1 | |
| | spearman | | 0.9698 | | | | 1 | | | 0.7913 | | | | 1 | |

**Table 3:** P-values from comparing methods from Figure 1c in the task of translation. The Mann–Whitney U test was used to compare the performance of the different methods

| | | A375 | | | | | | | | | HT29 | | | | | | | | |
|---|---|---|---|---|---|---|---|---|---|---|---|---|---|---|---|---|---|---|---|
| | | Autoencoders with classifier | CPA-based Autoencoders | DCS | DCS modified v1 | DCS modified v2 | FIT | TransCompR | direct translation | similarity-trained Autoencoders | Autoencoders with classifier | CPA-based Autoencoders | DCS | DCS modified v1 | DCS modified v2 | FIT | TransCompR | direct translation | similarity-trained Autoencoders |
| Autoencoders with classifier | pearson | 1 | 0.3447 | | | | 0.0539 | 0.9698 | | | 1 | 0.5205 | | | | 0.0028 | 0.6232 | | |
| | sign accuracy | 1 | 0.273 | | | | $\leq 10^{-3}$ | 0.8501 | | | 1 | 0.5205 | | | | $\leq 10^{-3}$ | 0.5205 | | |
| | spearman | 1 | 0.2123 | | | | $\leq 10^{-3}$ | 0.9698 | | | 1 | 0.6232 | | | | $\leq 10^{-3}$ | 0.6232 | | |
| CPA-based Autoencoders | pearson | | 1 | | | | 0.1405 | 0.3847 | | | | 1 | | | | | | | |
| | sign accuracy | | 1 | | | | $\leq 10^{-3}$ | 0.3447 | | | | 1 | | | | | | | |
| | spearman | | 1 | | | | $\leq 10^{-3}$ | 0.3447 | | | | 1 | | | | | | | |
| DCS | pearson | 0.273 | 0.6776 | 1 | 0.8501 | 0.0757 | 0.5205 | 0.3847 | 0.0312 | 0.3847 | 0.089 | 0.1859 | 1 | 0.9097 | 0.0211 | 0.5205 | 0.1405 | 0.0211 | 0.1859 |
| | sign accuracy | 0.3447 | 0.9698 | 1 | 0.9698 | 0.089 | $\leq 10^{-3}$ | 0.273 | 0.0257 | 0.6232 | 0.064 | 0.1859 | 1 | 0.9097 | 0.0539 | $\leq 10^{-3}$ | 0.1212 | 0.0173 | 0.3075 |
| | spearman | 0.4727 | 0.9698 | 1 | 1 | 0.089 | $\leq 10^{-3}$ | 0.3447 | 0.0257 | 0.6232 | 0.0757 | 0.273 | 1 | 0.9698 | 0.0539 | $\leq 10^{-3}$ | 0.1041 | 0.0113 | 0.2123 |
| DCS modified v1 | pearson | 0.3447 | 0.8501 | | 1 | 0.1041 | 0.5205 | 0.3847 | 0.0312 | 0.5205 | 0.089 | 0.1859 | | 1 | 0.0376 | 0.4274 | 0.1859 | 0.0211 | 0.1859 |
| | sign accuracy | 0.3447 | 0.9698 | | 1 | 0.089 | $\leq 10^{-3}$ | 0.273 | 0.0257 | 0.5708 | 0.064 | 0.273 | | 1 | 0.0376 | $\leq 10^{-3}$ | 0.1405 | 0.0173 | 0.273 |
| | spearman | 0.3447 | 1 | | 1 | 0.1041 | $\leq 10^{-3}$ | 0.3075 | 0.0257 | 0.6776 | 0.064 | 0.3075 | | 1 | 0.0376 | $\leq 10^{-3}$ | 0.1405 | 0.0113 | 0.2123 |
| DCS modified v2 | pearson | 0.4727 | 0.1041 | | | 1 | 0.0312 | 0.3075 | 0.0013 | 0.2413 | 0.4727 | 0.3075 | | | 1 | 0.0017 | 0.3075 | $\leq 10^{-3}$ | 0.4274 |
| | sign accuracy | 0.3075 | 0.1212 | | | 1 | $\leq 10^{-3}$ | 0.4727 | 0.0017 | 0.3447 | 0.5708 | 0.273 | | | 1 | $\leq 10^{-3}$ | 0.2413 | $\leq 10^{-3}$ | 0.3075 |
| | spearman | 0.273 | 0.1041 | | | 1 | $\leq 10^{-3}$ | 0.4274 | 0.001 | 0.2123 | 0.6776 | 0.3075 | | | 1 | $\leq 10^{-3}$ | 0.2413 | $\leq 10^{-3}$ | 0.273 |
| direct translation | pearson | 0.0022 | 0.0073 | | | | 0.0312 | 0.0058 | 1 | | $\leq 10^{-3}$ | $\leq 10^{-3}$ | | | | 0.014 | $\leq 10^{-3}$ | 1 | |
| | sign accuracy | 0.0022 | 0.0073 | | | | $\leq 10^{-3}$ | 0.0028 | 1 | | $\leq 10^{-3}$ | 0.001 | | | | $\leq 10^{-3}$ | $\leq 10^{-3}$ | 1 | |
| | spearman | 0.0022 | 0.0073 | | | | $\leq 10^{-3}$ | 0.0022 | 1 | | $\leq 10^{-3}$ | $\leq 10^{-3}$ | | | | $\leq 10^{-3}$ | $\leq 10^{-3}$ | 1 | |
| FIT | pearson | | | | | | 1 | 0.0757 | | | | 0.0257 | | | | 1 | 0.0091 | | |
| | sign accuracy | | | | | | 1 | $\leq 10^{-3}$ | | | | $\leq 10^{-3}$ | | | | 1 | $\leq 10^{-3}$ | | |
| | spearman | | | | | | 1 | $\leq 10^{-3}$ | | | | $\leq 10^{-3}$ | | | | 1 | $\leq 10^{-3}$ | | |
| similarity-trained Autoencoders | pearson | 0.5205 | 0.6232 | | | | 0.1041 | 0.7337 | 0.0036 | 1 | 0.5205 | 0.7913 | | | | 0.0173 | 0.9097 | $\leq 10^{-3}$ | 1 |
| | sign accuracy | 0.5708 | 0.7913 | | | | $\leq 10^{-3}$ | 0.5205 | 0.0046 | 1 | 0.5708 | 1 | | | | $\leq 10^{-3}$ | 1 | 0.001 | 1 |
| | spearman | 0.5708 | 0.8501 | | | | $\leq 10^{-3}$ | 0.7337 | 0.0036 | 1 | 0.6232 | 0.9698 | | | | $\leq 10^{-3}$ | 0.9097 | $\leq 10^{-3}$ | 1 |
| TransCompR | pearson | | | | | | | 1 | | | | 0.9698 | | | | | 1 | | |
| | sign accuracy | | | | | | | 1 | | | | 0.9698 | | | | | 1 | | |
| | spearman | | | | | | | 1 | | | | 0.7913 | | | | | 1 | | |

**Table 4:** P-values from comparing methods from Figure 1c in the task of reconstruction. The Mann–Whitney U test was used to compare the performance of the different methods

| | | A375 | | | | | | | HT29 | | | | | | |
|---|---|---|---|---|---|---|---|---|---|---|---|---|---|---|---|
| | | Autoencoders with classifier | CPA-based Autoencoders | DCS | DCS modified v1 | DCS modified v2 | TransCompR | similarity-trained Autoencoders | Autoencoders with classifier | CPA-based Autoencoders | DCS | DCS modified v1 | DCS modified v2 | TransCompR | similarity-trained Autoencoders |
| Autoencoders with classifier | pearson | 1 | 0.6776 | | | | $\leq 10^{-3}$ | | 1 | 0.8501 | | | | 0.0013 | |
| | sign accuracy | 1 | 0.6776 | | | | $\leq 10^{-3}$ | | 1 | 0.7337 | | | | 0.014 | |
| | spearman | 1 | 0.7337 | | | | $\leq 10^{-3}$ | | 1 | 0.7913 | | | | 0.0073 | |
| CPA-based Autoencoders | pearson | | 1 | | | | | | | 1 | | | | | |
| | sign accuracy | | 1 | | | | | | | 1 | | | | | |
| | spearman | | 1 | | | | | | | 1 | | | | | |
| DCS | pearson | $\leq 10^{-3}$ | $\leq 10^{-3}$ | 1 | $\leq 10^{-3}$ | $\leq 10^{-3}$ | $\leq 10^{-3}$ | $\leq 10^{-3}$ | $\leq 10^{-3}$ | $\leq 10^{-3}$ | 1 | $\leq 10^{-3}$ | $\leq 10^{-3}$ | $\leq 10^{-3}$ | $\leq 10^{-3}$ |
| | sign accuracy | $\leq 10^{-3}$ | $\leq 10^{-3}$ | 1 | $\leq 10^{-3}$ | $\leq 10^{-3}$ | $\leq 10^{-3}$ | $\leq 10^{-3}$ | $\leq 10^{-3}$ | $\leq 10^{-3}$ | 1 | $\leq 10^{-3}$ | $\leq 10^{-3}$ | $\leq 10^{-3}$ | $\leq 10^{-3}$ |
| | spearman | $\leq 10^{-3}$ | $\leq 10^{-3}$ | 1 | $\leq 10^{-3}$ | $\leq 10^{-3}$ | $\leq 10^{-3}$ | $\leq 10^{-3}$ | $\leq 10^{-3}$ | $\leq 10^{-3}$ | 1 | $\leq 10^{-3}$ | $\leq 10^{-3}$ | $\leq 10^{-3}$ | $\leq 10^{-3}$ |
| DCS modified v1 | pearson | $\leq 10^{-3}$ | $\leq 10^{-3}$ | | 1 | $\leq 10^{-3}$ | $\leq 10^{-3}$ | $\leq 10^{-3}$ | $\leq 10^{-3}$ | $\leq 10^{-3}$ | | 1 | $\leq 10^{-3}$ | $\leq 10^{-3}$ | $\leq 10^{-3}$ |
| | sign accuracy | $\leq 10^{-3}$ | $\leq 10^{-3}$ | | 1 | $\leq 10^{-3}$ | $\leq 10^{-3}$ | $\leq 10^{-3}$ | $\leq 10^{-3}$ | $\leq 10^{-3}$ | | 1 | $\leq 10^{-3}$ | $\leq 10^{-3}$ | $\leq 10^{-3}$ |
| | spearman | $\leq 10^{-3}$ | $\leq 10^{-3}$ | | 1 | $\leq 10^{-3}$ | $\leq 10^{-3}$ | $\leq 10^{-3}$ | $\leq 10^{-3}$ | $\leq 10^{-3}$ | | 1 | $\leq 10^{-3}$ | $\leq 10^{-3}$ | $\leq 10^{-3}$ |
| DCS modified v2 | pearson | $\leq 10^{-3}$ | $\leq 10^{-3}$ | | | 1 | $\leq 10^{-3}$ | $\leq 10^{-3}$ | $\leq 10^{-3}$ | $\leq 10^{-3}$ | | | 1 | $\leq 10^{-3}$ | $\leq 10^{-3}$ |
| | sign accuracy | $\leq 10^{-3}$ | $\leq 10^{-3}$ | | | 1 | $\leq 10^{-3}$ | $\leq 10^{-3}$ | $\leq 10^{-3}$ | $\leq 10^{-3}$ | | | 1 | $\leq 10^{-3}$ | 0.0017 |
| | spearman | $<10^{-3}$ | $<10^{-3}$ | | | 1 | $<10^{-3}$ | $<10^{-3}$ | $<10^{-3}$ | $<10^{-3}$ | | | 1 | $<10^{-3}$ | $<10^{-3}$ |
| similarity-trained Autoencoders | pearson | 0.7337 | 0.9097 | | | | $\leq 10^{-3}$ | 1 | 0.6776 | 0.9097 | | | | 0.0073 | 1 |
| | sign accuracy | 0.7337 | 0.6776 | | | | $\leq 10^{-3}$ | 1 | 0.9097 | 0.7337 | | | | 0.0312 | 1 |
| | spearman | 0.6776 | 0.5205 | | | | $\leq 10^{-3}$ | 1 | 0.7913 | 0.8501 | | | | 0.014 | 1 |
| TransCompR | pearson | | $\leq 10^{-3}$ | | | | 1 | | | 0.0058 | | | | 1 | |
| | sign accuracy | | $\leq 10^{-3}$ | | | | 1 | | | 0.0211 | | | | 1 | |
| | spearman | | $\leq 10^{-3}$ | | | | 1 | | | 0.0046 | | | | 1 | |

**Table 5:** Regularization λ terms in the loss function of different cases

| Case | $\lambda_{recon}$ | $\lambda_{distance}$ | $\lambda_{cosine}$ | $\lambda_{MI}$ | $\lambda_{prior}$ | $\lambda_{enc,i}$ | $\lambda_{dec,i}$ | $\lambda_{L2class,i}$ | $\lambda_{class,i}$ | $\lambda_{adverse}$ | $\lambda_{trained_{effect}}$ | $\lambda_{intermediate,enc}$ |
|---|---|---|---|---|---|---|---|---|---|---|---|---|
| L1000 10k genes | 1 | 1 | 40 | 100 | 1 | $10^{-2}$ | $10^{-2}$ | $10^{-2}$ | 500 | 500 | $10^{-4}$ | $10^{-5}$ |
| L1000 landmark genes | 1 | 10 | 10 | 100 | 1 | $10^{-2}$ | $10^{-2}$ | $10^{-2}$ | 1000 | 1000 | $10^{-4}$ | $10^{-5}$ |
| Lung fibrosis | 1 | 100 | 100 | 1 | 1 | $10^{-7}$ | $10^{-7}$ | $10^{-7}$ | 1 | 10 | $10^{-6}$ | $10^{-5}$ |
| Serology dataset | 1 | 40 | 70 | 100 | 1 | $10^{-6}$ | $10^{-6}$ | $10^{-4} - 10^{-5}$ | 100 | 100 | $10^{-4}$ | $10^{-5}$ |
